# Connecting models, networks and experiments: Revisiting the role of viruses in marine carbon cycling

**DOI:** 10.64898/2026.02.12.705374

**Authors:** Maxine Pruvôt, Bart Haegeman, Fabien Joux, David Demory

## Abstract

The marine microbial food web is a key driver of global carbon cycling, mediating the production, consumption and remineralisation of organic matter in the ocean. Within this food web, viruses exert strong top-down control by infecting and lysing both phytoplankton and heterotrophic bacteria. Although virus-induced mortality is recognised as a factor shaping microbial dynamics, accurately quantifying its contribution to carbon cycling remains difficult. A range of approaches have been used to estimate this contribution, notably: (1) dynamical microbial ecosystem models, (2) static carbon flux networks, and (3) direct measurements of virus-induced mortality, such as dilution assays. In this study, we bring these approaches together by formulating them within a common mathematical framework. This enables us to clarify the difference between two notions of lysis percentage in the literature, to compare predictions of previous microbial ecosystem models and flux network reconstructions, and to critically evaluate the theoretical approach of eliminating viruses from the system to explore their effects. Our analysis clarifies how the different approaches capture complementary aspects of the role of marine viruses in carbon cycling across varying spatial and temporal scales.

## 1 Introduction

The ocean plays a central role in regulating atmospheric carbon through a combination of physical, chemical and biological processes. Central to this regulation is the biological processing of carbon in the upper ocean (Boyd *et al*., 2019). In surface waters, phytoplankton fix inorganic CO_2_ into organic carbon through photosynthesis. The fate of this newly produced organic matter is complex, involving multiple pathways. For simplicity, two main routes can be distinguished (Azam *et al*., 1983). First, organic carbon can be transferred to higher trophic levels as phytoplankton are grazed by zooplankton. Second, it can enter the microbial loop through the release of dissolved organic matter that fuels the growth of heterotrophic bacteria that remineralise organic into inorganic matter. Through these interconnected processes, carbon is continually transformed, respired and recycled within the ocean’s upper layers, driven by microbial growth and mortality (Falkowski *et al*., 2008).

Among the factors controlling microbial mortality, viruses are particularly influential. They are the most abundant biological entities in the sea and, lacking their own metabolism, replicate by infecting living cells (Suttle, 2007). Viruses that infect phytoplankton directly affect primary producers (Suttle *et al*., 1990), while viruses targeting heterotrophic bacteria, called bacteriophages, impact the recycling of organic matter through the microbial loop (Weinbauer & Peduzzi, 1994, Proctor & Fuhrman, 1990). When lytic infection occurs, the host cell bursts, releasing its contents and viral particles into the environment. This release of cellular material, a process known as the viral shunt, redirects organic carbon away from higher trophic levels and back into the dissolved pool (Wilhelm & Suttle, 1999). In doing so, viral lysis fuels microbial activity and enhances carbon recycling within the microbial loop, thereby reshaping the pathways of carbon flow in the surface ocean (Fuhrman, 1999, Middelboe & Lyck, 2002, Suttle, 2005, Wei *et al*., 2025).

A quantitative understanding of viral impacts is essential for studying carbon cycling in the ocean. In simplified descriptions of the microbial food web, viral losses are often lumped together with other forms of mortality, such as grazing (Moore *et al*., 2004, Follows *et al*., 2007, Aumont *et al*., 2015). This may be adequate for a static view of carbon fluxes, but it lacks the mechanistic detail needed to understand the dynamics of carbon processing. For instance, distinguishing the fraction of mortality caused by viruses is important for assessing how the system responds to environmental perturbations, such as warming (Demory *et al*., 2021, Krishna *et al*., 2024) or nutrient shifts (Thingstad, 2000, McDaniel & Paul, 2005, Maat & Brussaard, 2016, Pourtois *et al*., 2020). Despite the importance of such quantification, measuring the role of viruses remains difficult. Viral infection and lysis occur at microscopic scales and over short timescales, making direct observation challenging (Vincent & Vardi, 2023). At larger spatial or temporal scales, measurements often integrate other sources of mortality. Adding to the complexity, viral effects propagate through trophic interactions and feedbacks, and are difficult to disentangle from other processes unrelated to viruses (Beckett *et al*., 2024, Talmy *et al*., 2025).

This complexity has been explored through a range of quantitative approaches. On-board experimental methods, such as dilution assays, are used to infer viral impacts on microbial dynamics (Baudoux *et al*., 2006, 2008, Kimmance & Brussaard, 2010, Evans & Brussaard, 2012, Mojica & Brussaard, 2026). At the other end of the spectrum, modelling approaches describe the dynamics of entire microbial ecosystems and use observations to constrain model parameters (Weitz *et al*., 2015). Between these lie carbon flux network reconstructions, which estimate carbon fluxes between microbial compartments while neglecting their dynamics (Fuhrman, 1999, Wilhelm & Suttle, 1999, Mojica, 2015). In this study, we bring structure to this diversity of methods by placing them on a common quantitative footing (Sect. 2). This allows us to highlight both connections and differences among approaches. We then illustrate its use through three applications: a conceptual clarification of lysis percentages, often employed to quantify viral impacts (Sect. 3); a comparison of previous modelling studies, revealing large divergences (Sect. 4); and a critical assessment of a common theoretical practice of eliminating viruses to evaluate their effects (Sect. 5).

## 2 Quantitative approaches to assess the role of viruses

In this section we describe the three quantitative approaches considered in this study: dynamical system models, carbon flux network reconstructions, and direct measurements of viral lysis. Their main assumptions and conceptual differences are illustrated in Fig. 1; main terms are summarised in Table 1.

**Table 1:** Glossary of terms in microbial ecosystem modelling.

| Term | Explanation |
| --- | --- |
| Dynamical system | A modelling approach based on a set of differential equations that describe how the variables of a system change over time. In the context of microbial ecosystems, these variables include the abundances of different microbial populations and the sizes of the carbon pools they produce and consume. |
| Dynamical variable | A quantity in a DS model that changes over time, such as the abundance of a microbial population or the size of a carbon pool. |
| Model parameter | A quantity in a DS model that is constant over time and determines the strength or rate of processes, such as growth, consumption or mortality. |
| Flux network | A modelling approach that represents the distribution of carbon fluxes among ecosystem compartments. These models focus on the flows themselves rather than on changes in compartment sizes over time. |
| Compartment | A component of a FN model, biotic or abiotic, that receives and releases fluxes of matter. |
| Partitioning ratio | In a FN model, the fraction of carbon leaving a source compartment that is transferred to a specific recipient compartment. |
| Flux ratio | A flux expressed as a proportion of the total primary production. FN models provide these relative rather than absolute fluxes. |
| Through-flow flux | In a FN model, sum of all incoming fluxes to a recipient compartment. |
| Turnover rate | The rate at which the stock of a compartment is renewed. Its inverse is called the turnover time. |
| Lysis process rate | The rate at which viral lysis occurs for phytoplankton or heterotrophic bacteria, with units of inverse time. It is often expressed as a percentage per day and is sometimes called the lysis percentage. |
| Lysis flux ratio | The total carbon flux from viral lysis of phytoplankton and heterotrophic bacteria relative to primary production. It is often expressed as a percentage, called the lysis percentage. |

**Figure 1.**
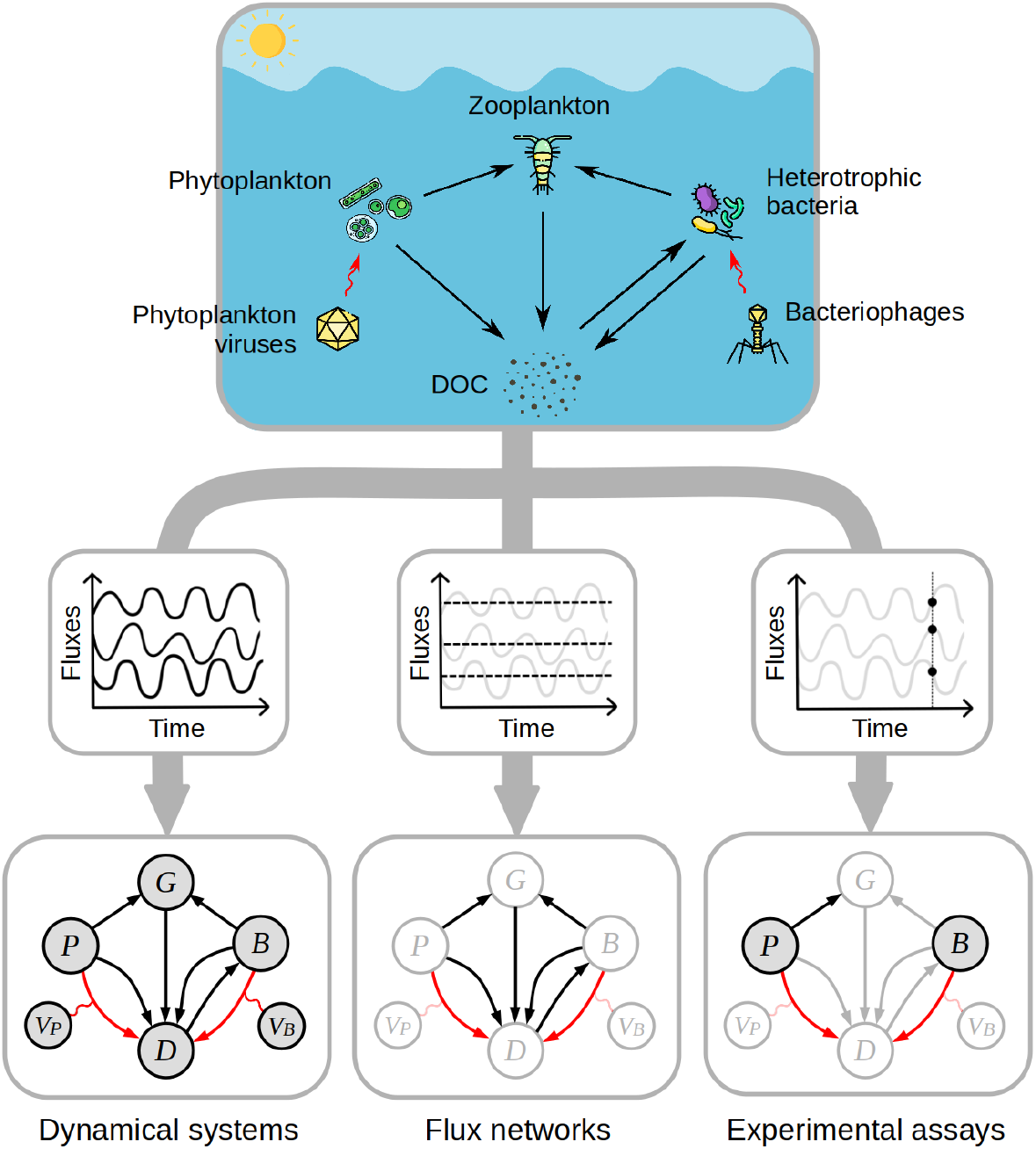
Contrasting quantitative approaches to study the role of marine viruses. We use a simplified representation of the marine microbial food web, in which a limited number of compartments and the carbon fluxes between them are considered. In a first approach (leftmost arrow), dynamical system models are used to describe the temporal evolution of carbon stocks and fluxes. This approach provides a comprehensive view of ecosystem dynamics, but it requires detailed empirical knowledge for model parameterisation. In a second approach (central arrow), carbon stocks are not explicitly represented. Instead, the system is described as a network of balanced carbon fluxes between compartments, forming a stationary flux network. This approach provides more limited insight into ecosystem functioning, but requires fewer parameters. In a third approach (rightmost arrow), ecosystem processes are quantified experimentally. In the context of marine viruses, the most common methods include the modified dilution assay and the virus reduction assay. These methods provide quantitative estimates of selected stocks and fluxes, but they are typically limited to snapshot observations and do not capture ecosystem dynamics. The virus-mediated fluxes are represented red arrows. The icons are made by Freepik from Flaticon.

### Dynamical system (DS) model

Dynamical system models describe how microbial populations and the carbon pools they produce and consume change over time. Models of this type vary widely in their level of complexity. For spatial structure, some represent vertical gradients in the water column or differences between ocean regions (Follows *et al*., 2007, Krishna *et al*., 2024), while others treat the system as spatially uniform. In terms of temporal structure, some resolve seasonal or daily variability (Beckett *et al*., 2024), whereas others represent temporally averaged conditions and thus assume a constant state over time (Weitz *et al*., 2015, Pourtois *et al*., 2020). Models also vary in how they represent biological diversity, ranging from aggregated community compartments to detailed taxonomic or functional group structure (Follows *et al*., 2007).

Here we focus on models that capture the essential processes in the ecosystem while remaining as simple as possible. A minimal version of such a model, detailed in the Supporting Information Sect. S1, includes phytoplankton *P*, heterotrophic bacteria *B*, their grazers *G*, and the viral populations *V*_*P*_ and *V*_*B*_, together with a pool of dissolved organic carbon *D* (see Table 2 for a summary of the mathematical notation; see also Supporting Information Table S2). Each process is then represented by a few parameters. For example, consider the infection of phytoplankton by their viruses. Phytoplankton cells become infected at a rate proportional to the product of host and virus abundances, characterised by the infection rate *ϕ*_*P*_ . Each lysed host releases on average *β*_*P*_ new viral particles, equal to the burst size. Viruses are also subject to decay and other loss processes, summarised by a loss rate 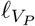 Together, these constant parameters determine the viral control of phytoplankton biomass and the corresponding release of organic carbon upon lysis.

**Table 2:** Overview of the mathematical notation used in this paper. A more detailed description is given in the appendices.

| Variables in DE models |  | units |
| --- | --- | --- |
| $P$ | phytoplankton abundance | cells L<sup>-1</sup> |
| $P_c$ | phytoplankton carbon stock, i.e., $P_c = q_P P$<br>with $q_P$ the carbon quota of phytoplankton cells | $\mu\text{mol of carbon L}^{-1}$ |
| $B$ | abundance of heterotrophic bacteria | cells L<sup>-1</sup> |
| $B_c$ | carbon stock of heterotrophic bacteria | $\mu\text{mol of carbon L}^{-1}$ |
| $V_P$ | abundance of phytoplankton viruses | viruses L<sup>-1</sup> |
| $V_B$ | abundance of bacteriophages | viruses L<sup>-1</sup> |
| $G$ | abundance of zooplankton grazers | individuals L<sup>-1</sup> |
| $D$ | concentration of dissolved organic carbon | $\mu\text{mol of carbon L}^{-1}$ |
| Parameters in DE models |  |  |
| $\phi_P$ | infection rate of phytoplankton viruses | L/virus d<sup>-1</sup> |
| $\beta_P$ | burst size of phytoplankton viruses | viruses/cell |
| $\ell_{V_P}$ | loss rate of phytoplankton viruses | d<sup>-1</sup> |
| ... | see Table S2 for the full list of model parameters |  |
| Fluxes in FN models |  |  |
| $F_{PG}$ | carbon flux from phytoplankton to grazer compartment | $\mu\text{mol of carbon L}^{-1} \text{ d}^{-1}$ |
| $F_{BD}^V$ | virus-mediated carbon flux from $B$ to $D$ compartment | $\mu\text{mol of carbon L}^{-1} \text{ d}^{-1}$ |
| $F_{BD}^0$ | virus-independent carbon flux from $B$ to $D$ compartment,<br>i.e., $F_{BD}^0 = F_{BD} - F_{BD}^V$ | $\mu\text{mol of carbon L}^{-1} \text{ d}^{-1}$ |
| $F_P^{\text{tot}}$ | through-flow flux of phytoplankton compartment,<br>i.e., primary production | $\mu\text{mol of carbon L}^{-1} \text{ d}^{-1}$ |
| $F_B^{\text{tot}}$ | through-flow flux of $B$ compartment | $\mu\text{mol of carbon L}^{-1} \text{ d}^{-1}$ |
| $f_{PG}$ | carbon flux from phytoplankton to grazer compartment,<br>relative to primary production, i.e., $f_{PG} = F_{PG}/F_P^{\text{tot}}$ | — |
| $f_{BD}^V$ | virus-mediated carbon flux from $B$ to $D$ compartment<br>relative to primary production, i.e., $f_{BD}^V = F_{BD}^V/F_P^{\text{tot}}$ | — |
| $f_{BD}^0$ | virus-independent carbon flux from $B$ to $D$ compartment<br>relative to primary production, i.e., $f_{BD}^0 = f_{BD} - f_{BD}^V$ | — |
| $f_B^{\text{tot}}$ | through-flow flux of $B$ compartment<br>relative to primary production, i.e., $f_B^{\text{tot}} = F_B^{\text{tot}}/F_P^{\text{tot}}$ | — |
| $a_{PG}$ | flux partitioning ratio from $P$ to $G$ compartment,<br>i.e., $a_{PG} = F_{PG}/F_P^{\text{tot}}$ | — |
| $a_{BD}^V$ | virus-mediated flux partitioning ratio from $B$ to $D$,<br>i.e., $a_{BD}^V = F_{BD}^V/F_B^{\text{tot}}$ | — |
| $a_{BD}^0$ | virus-independent flux partitioning ratio from $B$ to $D$,<br>i.e., $a_{BD}^0 = F_{BD}^0/F_B^{\text{tot}}$ | — |

Table 2 (*continued*)
| Lysis percentages |  | units |
| --- | --- | --- |
| $\mathcal{L}_P^{\text{proc}}$ | rate of phytoplankton loss due to viral lysis<br>i.e., $\mathcal{L}_P^{\text{proc}} = F_{PD}^V / P_c$ | $\text{d}^{-1}$ or $\% \text{ d}^{-1}$ |
| $\mathcal{L}_B^{\text{proc}}$ | rate of heterotrophic bacteria loss due to viral lysis<br>i.e., $\mathcal{L}_B^{\text{proc}} = F_{BD}^V / B_c$ | $\text{d}^{-1}$ or $\% \text{ d}^{-1}$ |
| $\mathcal{L}^{\text{flux}}$ | viral lysis flux ratio<br>i.e., $\mathcal{L}^{\text{flux}} = (F_{PD}^V + F_{BD}^V) / F_P^{\text{tot}} = f_{PD}^V + f_{BD}^V$ | — or $\%$ |

In practice, setting up such models involves substantial uncertainty because most parameters are poorly constrained. Reported values in the literature often span several orders of magnitude, reflecting real-world variability, but also differences in experimental conditions, species composition or measurement methods. To account for this, models can be explored across broad, plausible parameter ranges, rather than relying on a single best estimate (Weitz *et al*., 2015). This approach captures a spectrum of possible model outcomes. By comparing these outcomes to empirical observations, one can obtain an ensemble of plausible parameter combinations.

This ensemble of parameter sets can be used to explore model predictions about the role of viruses. This can be done by examining virus-related processes, such as the fraction of phytoplankton mortality caused by viral lysis (Beckett *et al*., 2024). Another approach is to compare the steady state of the full system, which includes viruses, with that of a system where viral populations are removed (Weitz *et al*., 2015, Pourtois *et al*., 2020). Differences in population abundances or carbon fluxes then provide a measure of how viral processes shape ecosystem structure and function. Overall, dynamical system models offer a flexible framework for investigating the mechanisms that govern the microbial ecosystem.

### Flux network (FN) model

Flux network reconstruction focuses on estimating the magnitude of carbon flows between compartments of the microbial ecosystem (Fuhrman, 1992, 1999, Wilhelm & Suttle, 1999). Compared with dynamical system models, which describe changes in compartment sizes over time, flux network models are more limited because they consider only the flows between compartments. They represent the system as a set of steady-state exchanges of carbon among components. This approach makes it possible to see how primary production is redistributed across microbial groups and how carbon is ultimately lost from the system through respiration or export.

In practice, the method relies on specifying partitioning ratios that determine how the outgoing carbon flux from a given compartment is divided among downstream compartments. For each compartment *i*, the coefficients *a*_*ij*_ describe the fraction of its total outgoing flux directed toward compartment *j* (see Supporting Information eq. S3). The collection of all partitioning ratios defines the structure of the network and allows the stationary flux distribution to be computed, up to a multiplicative constant (see Supporting Information Sect. S2 for details). Typically, all fluxes are expressed relative to primary production, which serves as the reference flux in the system.

An important distinction between flux network analysis and DS models is that the former provides no direct information about time scales. To infer these, one must consider, in addition to the flux distribution, the standing stocks of the compartments. Explicitly, the turnover rate of a compartment is obtained as its through-flow flux, which is the total flux entering (and equivalently leaving) that compartment, divided by its standing stock (see Supporting Information Sect. S2). Hence, flux network analysis provides stationary fluxes, but not their dynamics, underlying stocks or turnover times.

A second difference from the DS modelling approach lies in the way partitioning ratios are treated. In flux network analysis, the partitioning ratios *a*_*ij*_ are assumed to be fixed and prescribed. In contrast, in DS models these ratios emerge from the system dynamics, as they depend on the equilibrium biomasses of the interacting populations. For example, in the case of phytoplankton, the outgoing carbon flux is partitioned among grazing by zooplankton, viral lysis and loss to respiration. In a mechanistic model, the relative magnitudes of these processes depend on the standing stocks of grazers, viruses and phytoplankton themselves, and thus vary with system state.

### Experimental measurement of viral lysis

Carbon fluxes associated with viral lysis can be measured using several experimental approaches. Among these, the modified dilution assay and the virus reduction assay are generally considered as the most quantitative methods.

### Modified dilution (MD) assay

The modified dilution assay (Evans *et al*., 2003, Baudoux *et al*., 2007, Kimmance & Brussaard, 2010) is based on the original dilution method of Landry & Hassett (1982). It generates a series of parallel dilutions by combining natural seawater with seawater selectively filtered to remove either grazers or viruses. The resulting samples are incubated under controlled conditions, and changes in phytoplankton abundance are monitored over time.

In an idealised theoretical setting (Beckett & Weitz, 2018; see Supporting Information Sect. S3), this assay enables estimation of *F*_*PG*_, the carbon flux from phytoplankton to grazers, and 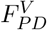, the flux from phytoplankton to dissolved organic carbon generated by viral lysis. By dividing these fluxes by the standing stock of phytoplankton carbon, one obtains the corresponding phytoplankton mortality rates due to grazing and viral infection, respectively. In experimental conditions, several processes can introduce deviations from this ideal behavior, for example, changes in resource concentrations, shifts in phytoplankton density during dilution, or the natural diversity of host-virus pairs, which contrasts with the single-pair assumption of the theoretical analysis.

### Virus reduction (VR) assay

The virus reduction assay (Wilhelm *et al*., 2002, Weinbauer *et al*., 2010, Lara *et al*., 2017) starts with a filtration step that strongly reduces free viruses while leaving host cells largely intact. The treated sample is subsequently incubated under conditions similar to those of the natural environment. During incubation, viruses that were already infecting host cells complete their replication cycle and are released into the surrounding water, allowing the accumulation of new free viruses to be measured over time.

Under idealised theoretical conditions (see Supporting Information Sect. S3), this assay allows estimation of the carbon flux 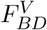 from bacteria to dissolved organic carbon caused by viral lysis, together with the corresponding virus-induced bacterial mortality rate. In experimental conditions, several factors can cause deviations from this ideal behavior, for example, uncertainty in burst size estimates, the presence of multiple host-virus pairs rather than a single representative one, or the exclusion of grazers during incubation, which may alter bacterial dynamics.

Together, these methods provide complementary insights into virus-mediated carbon transformations in the ocean. MD and VR experiments focus on short-term dynamics, quantifying the immediate impacts of viral infection on host populations. FN models capture long-term balances between compartments, averaging out spatial and temporal variation. DS models offer a more flexible framework that in principle could link different time scales. However, this would require detailed and comprehensive data, which are rarely available.

## 3 Lysis percentages: process rates vs flux ratios

The term lysis percentage is widely used to quantify the impact of viruses in microbial ecosystems. However, it can refer to two distinct quantities that are not directly related. In this section we clarify the difference between these two notions.

The first notion is based on process rates such as those obtained from modified dilution (MD) and virus reduction (VR) assays. These represent the rate at which host biomass is lost due to viral lysis and are defined as:

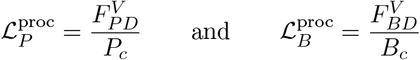

for phytoplankton and bacteria, respectively. It is common to express the rates 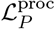 and 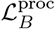 as lysis percentages. For example, if an MD assay yields 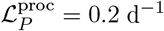, this is reported as “viral lysis removes 20% of the phytoplankton population per day.” Note that this percentage exceeds 100% if the lysis rate is faster than one per day.

The second notion arises from flux network (FN) analyses. Because these analyses provide only relative fluxes, results are commonly expressed relative to primary production 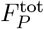. For example, FN models do not predict the absolute viral lysis flux 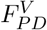 (e.g., in *µ*mol-C L^−1^ d^−1^), but instead estimate the relative flux 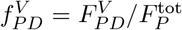, which is dimensionless. The ratio of total lysis flux to primary production,

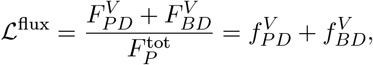

is then interpreted as a measure of the contribution of viral lysis to carbon cycling. The ratio ℒ^flux^ is also commonly expressed as a lysis percentage. For example, one may encounter statements such as “30% of primary production passes through the viral shunt,” corresponding to ℒ^flux^ = 0.3. Fluxes used in the FN approach should not be interpreted as a proportion of primary production, but instead as fluxes expressed in units of primary production. As we will see shortly, this percentage can also exceed 100%, but for reasons quite different from those for the lysis process rates.

The lysis process rates 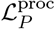 and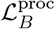 and the lysis flux ratio ℒ^flux^ cannot be compared, simply because they do not have the same dimensions. Process rates have dimensions of reciprocal time, while flux ratios are dimensionless. To illustrate this further, consider the case of phytoplankton viral lysis. Suppose that the phytoplankton lysis process rate is fixed at 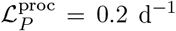. This same rate may occur under very different conditions. In a highly productive environment, with phytoplankton turnover rate *µ*_*P*_ = 2.0 d^−1^, the lysis flux ratio is ℒ^flux^ = 10%. In a less productive environment, where *µ*_*P*_ = 0.25 d^−1^, the lysis flux ratio is ℒ^flux^ = 80%. Thus, identical lysis process rates can yield very different lysis flux ratios, depending on the productivity of the system.

The decoupling between lysis process rates and lysis flux ratio can also occur in more subtle ways. For example, carbon released through viral lysis can be recycled within the microbial loop and pass again through additional viral lysis events. This recycling increases the lysis flux, and hence the lysis flux ratio ℒ^flux^, without necessarily changing the underlying lysis process rate. To illustrate this effect, we consider three FN models that share the same lysis partitioning ratios 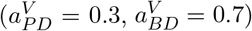, but differ in the strength of DOC recycling (Fig. 2; see Supporting Information Sect. S4 for details). The resulting lysis flux ratios are ℒ^flux^ = 35% for weak recycling, ℒ^flux^ = 58% for strong recycling, and ℒ^flux^ = 108% for very strong recycling. This shows that DOC recycling can substantially increase ℒ^flux^ and, consequently, the apparent importance of viral lysis for carbon cycling. Note that ℒ^flux^ larger than 100% can arise when the same carbon is cycled multiple times through the viral shunt.

**Figure 2.**
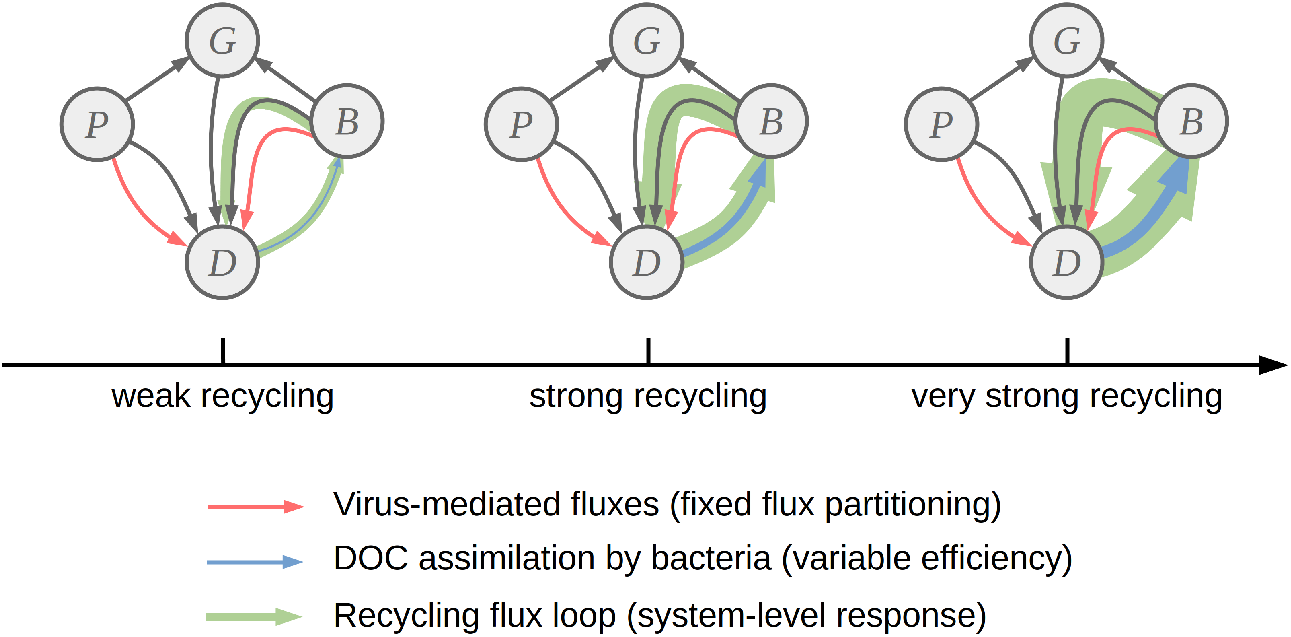
Conceptual illustration of the effect of DOC recycling on the viral lysis flux. Three flux networks are shown that differ in the efficiency of bacterial assimilation of DOC. This efficiency controls the strength of DOC recycling through the microbial loop. Enhanced bacterial assimilation increases system-level recycling, leading to larger lysis flux ratio ℒ^flux^, despite identical virus-mediated flux partitioning in the three networks. Flux magnitudes are illustrative; quantitative details are provided in Supporting Information Sect. S4.

In summary, the two notions of lysis percentage capture different aspects of viral impact. The process-based lysis percentages, derived from MD and VR assays, describe short-term population dynamics, quantifying how rapidly viruses remove hosts under given conditions. In contrast, the flux-based lysis percentage reflects the long-term role of viruses in shaping carbon pathways across the ecosystem.

## 4 Common ground to compare modelling approaches

Both dynamical system (DS) and flux network (FN) approaches have been used to quantify virus-mediated carbon fluxes in marine systems, but their results have not previously been compared. Here, we provide such a comparison across a selection of published studies (see Table 3).

**Table 3:** List of models used in PCA analysis. Model codes consist of the first three letters of the first author’s surname followed by a short code specifying the model variant, such as region or process type.

| Code | Type | Comments | Reference |
| --- | --- | --- | --- |
| FUH-B,<br>FUH-BP | FN | Foundational FN model. FUH-B includes only bacteria-infecting viruses, while FUH-BP includes both bacteria- and phytoplankton-infecting viruses. | Fuhrman (1992, 1999) |
| WIL | FN | Closely related to FUH models, but accounting for parameter uncertainty by using ranges of flux partitioning ratios rather than single values. | Wilhelm & Suttle (1999) |
| WEI | DS | Microbial ecosystem model without spatial structure, with model outcome uncertainty explored using ranges of parameter values. | Weitz *et al.* (2015) |
| MOJ-S,<br>MOJ-N | FN | Model parameterised with experimental assay data from two regional sites in the Northeast Atlantic: south (MOJ-S) and north (MOJ-N). | Mojica (2015) |
| XIE-H,<br>XIE-A | DS | Vertically resolved microbial ecosystem model extended to include viruses and parameterised with observational data from the Hawaii Ocean Time-series (XIE-H) and the Arabian Sea (XIE-A). | Xie *et al.* (2022) |

We first defined a common flux-network structure, specifying only the compartments and allowable carbon transfers, without assigning numerical values to the fluxes. We then projected each selected model (both DS and FN), for which both structure and parameters are fully specified, onto this reference framework (see Supporting Information Sect. S5). This mapping allowed us to extract through-flows and partitioning ratios from each model in a consistent way, enabling a consistent comparison across different models.

Fig. 3A presents the results of a principal component analysis (PCA) performed on the extracted variables. The first principal component (PC1, accounting for 84% of total variance) represents a gradient from grazing- to virus-driven phytoplankton loss. The phytoplankton lysis ratio 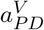 and the total DOC flux from viral lysis 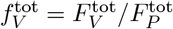 are positively correlated with PC1, whereas the phytoplankton grazing ratio *a*_*P G*_ and grazer through-flow flux 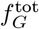 are negatively correlated with PC1.

**Figure 3.**
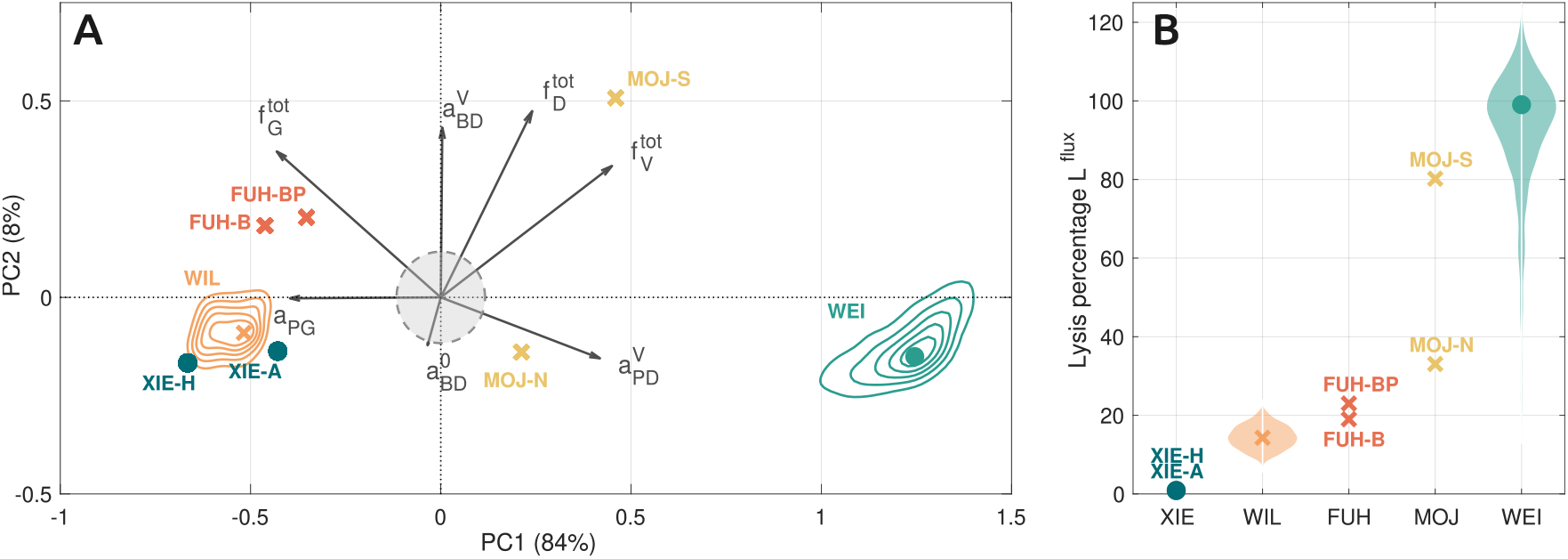
Previously published models on the role of marine viruses give very different results. We mapped all the models of Table 3 on a common reference model and used the variables of this reference model for further analysis. Panel A: Principal component analysis (PCA) of the models. Individual models are shown in color; circles denote DS models and crosses denote FN models. The variability within the WIL and WEI model sets is represented by density contours. Variable loadings are shown as grey arrows. Only the most influential loadings are displayed; less influential variables are located within the grey circle centered at the origin of the PCA plane. See Supporting Information Sect. S5 for the full list of variables. Panel B: Viral lysis flux ratio expressed as a percentage of primary production 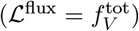 for the different models. The variability within the WIL and WEI model sets is shown using violin plots.

Model positions along PC1 reflect this gradient. The model realisations of WEI show strongly positive values of PC1, indicating that phytoplankton mortality is primarily driven by viral lysis. The models of MOJ have positive but smaller PC1 values, corresponding to ecosystems with intermediate contributions from viral lysis. In contrast, WIL, FUH and XIE exhibit strongly negative PC1 values, indicating ecosystems where phytoplankton mortality is mainly driven by grazing.

The second principal component (PC2, explaining 8% of total variance) captures smaller-scale variation. It is correlated with several variables, including the viral lysis ratio of heterotrophic bacteria 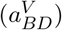. Overall, the PCA indicates that viral and grazing processes are important sources of variability across the compared studies.

Fig. 3B shows the distribution of viral lysis flux ratios, defined as the viral lysis flux expressed as a percentage of primary production. Across all models, these ratios span a wide range, from close to 0% to well above 100%. Values exceeding 100% arise only in specific flux networks that allow repeated recycling of carbon through viral lysis. Therefore, the models cover nearly the full range of allowed viral lysis flux ratios. This indicates substantial uncertainty in current model-based estimates of viral contributions to carbon cycling.

It is worth noting that the total spread among studies is also present within each modelling approach. Both the DS and FN subsets display a comparable range of variability in their predicted virus-mediated carbon fluxes. This indicates that the observed differences arise primarily from differences in parameter values rather than from the modelling approach itself.

## 5 Virus removal as a tool to quantify their role

Rather than directly quantifying virus-mediated fluxes, the impact of viruses can also be evaluated by comparing model configurations with and without viral components (Fuhrman, 1992, 1999, Wilhelm & Suttle, 1999, Weitz *et al*., 2015, Pourtois *et al*., 2020, Xie *et al*., 2022). In this section, we explore the type of information that can be extracted from such virus-removal analyses.

### Predicting virus removal effects from viral lysis fluxes

A first question we address is whether viral lysis fluxes can predict the effects of virus removal. To explore this, we examined correlations across model realisations between lysis fluxes and the changes observed after removing viruses in quantities such as primary production or microbial loop fluxes (see Supporting Information Sect. S6).

When we applied this analysis to an existing DS model (Weitz *et al*., 2015), removing viruses had drastic effects. Entire ecosystem compartments systematically collapsed, leading to a destructured system. This outcome can be seen as evidence that viruses maintain ecosystem persistence, but it also indicates that the model lacks processes allowing the ecosystem to remain functional in their absence.

We therefore constructed a simpler model in which this problem was not structurally present. The model implements the structure of the ecosystem represented in Fig. 1; we provide a detailed model description in Supporting Information Sect. S1. In 13% of the realisations, virus removal led to a destructured ecosystem; these cases were excluded from further analysis.

We then studied, for the 8651 remaining model realisations, the correlations between, on the one hand, variables in the model with viruses that are directly related to viral lysis and, on the other hand, the response of general ecosystem variables to virus removal. Fig. 4 illustrates the correlation patterns for a few representative pairs of variables (similar patterns were observed for other variable pairs, see Fig. S3). As measures of viral lysis, we considered the total lysis flux 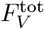 and the total lysis flux relative to primary production 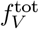. As response variables to virus removal, we considered the absolute and relative change in primary production,

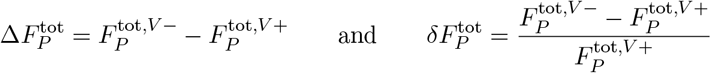

where 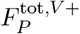 and 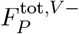 denote primary production with and without viruses, respectively.

**Figure 4.**
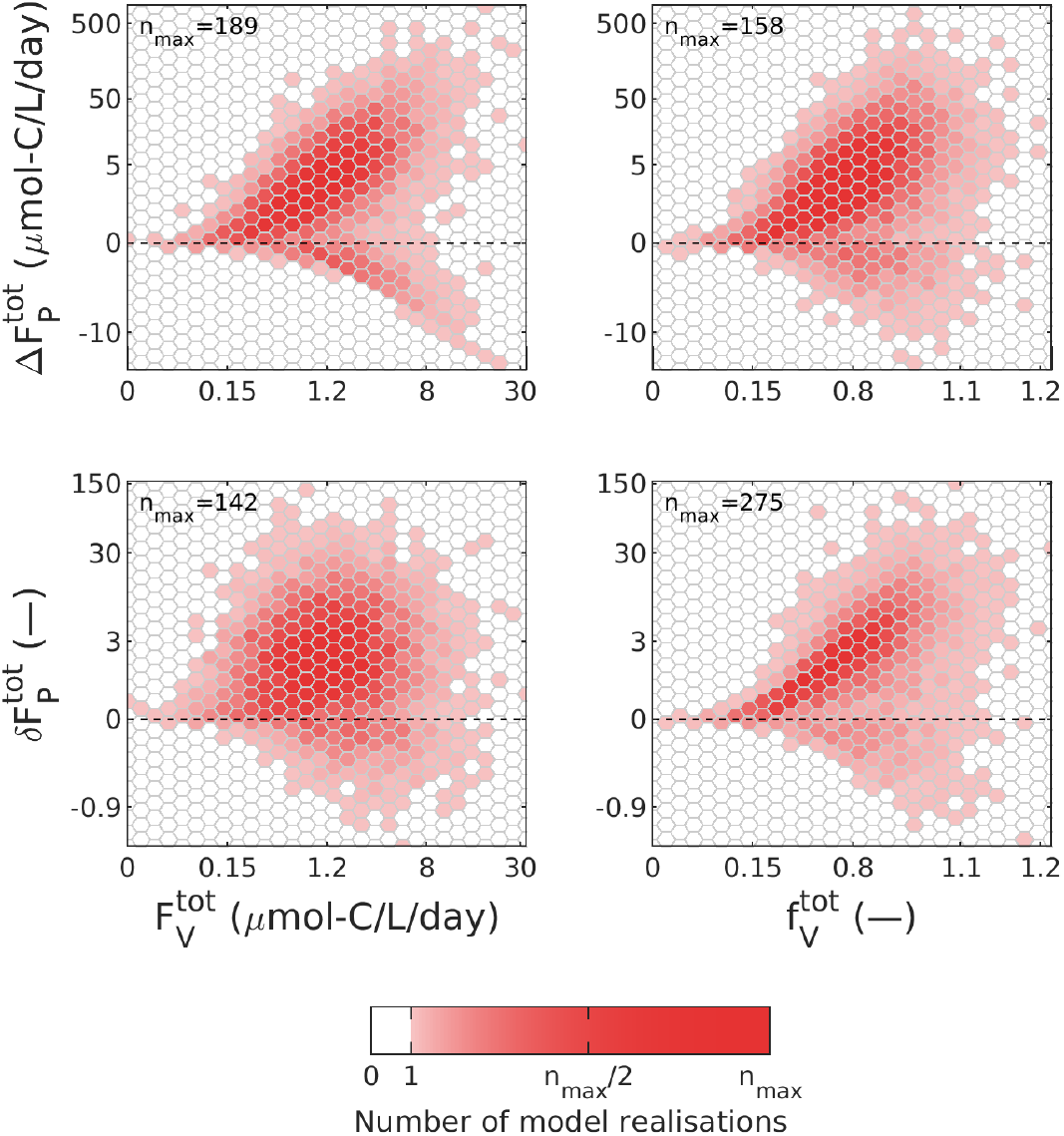
Do viral lysis fluxes predict the system’s response to virus removal? We studied the effects of viral removal using a simple microbial ecosystem model. Each panel shows the relationship between a viral lysis flux (*x*-axis) and the change of an ecosystem flux induced by virus removal (*y*-axis). For the *x*-axis: the absolute viral lysis flux 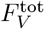 in the left panels; the viral lysis flux 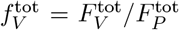 relative to primary production in the right panels. For the *y*-axis: the absolute change in primary production 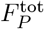 in the top panels; the relative change in 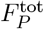 in the bottom panels. Each variable was transformed so that its values follow a Gaussian distribution while preserving their relative ordering. Colors indicate the number of model realisations in each hexagonal bin, normalised by the maximum bin count *n*_max_, which is indicated in each panel.

When viral lysis fluxes are small, the ecosystem shows only minor responses to virus removal. However, when viral lysis represents a substantial component of the flux network, ecosystem responses vary widely, ranging from an order-of-magnitude decrease to a two-orders-of-magnitude increase in primary production. Overall, correlations between viral lysis fluxes and ecosystem responses to virus removal are weak. A stronger correlation is observed for the pair 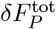 versus 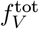 (bottom-right panel), but this relationship is partly artificial, as both variables share primary production in the virus-including system as a common normalisation factor.

We conclude that when viruses make a major contribution to carbon flows, the consequences of their removal are difficult to predict. This is not surprising: removing viruses leads to a reorganisation of carbon pathways. The viral lysis flux itself contains little information about how this reorganisation unfolds, and thus cannot predict the direction or magnitude of the resulting ecosystem response.

### Contrasting virus removal effects in DS and FN models

A second question we address is whether the effects of virus removal are consistent between dynamical system (DS) and flux network (FN) models. To investigate this, we generated model realisations of a simplified version of the DS model from Weitz *et al*. (2015), from which we constructed the corresponding FN model, and compared the impacts of virus removal in both model types (see Supporting Information Sect. S6).

As described above, we observed that in some model realisations (13% of the total set), virus removal in the DS model led to the extinction of one or more compartments. This outcome does not occur in the FN models, because compartment stocks are not explicitly represented. This observation highlights a fundamental divergence between DS and FN models in their responses to virus removal. We excluded model realisations in which virus removal caused population extinction in the DS model from further analysis.

Fig. 5 illustrates the extent to which the simpler FN approach can reproduce the effects of virus removal observed in the DS model. We consider several through-flow fluxes and partitioning ratios. Some fluxes, such as the through-flow of the bacterial compartment, are reproduced reasonably well by the FN approach, indicating that certain aspects of system functioning are captured consistently between the two model types. However, for most fluxes, we observe strong and systematic deviations. For instance, the through-flow flux of the grazer compartment is consistently lower in the FN models than in the DS models. This difference could arise because, in the DS approach, virus removal can increase the grazer population through higher host availability, leading to a larger inflow from the host compartments (phytoplankton and heterotrophic bacteria). This feedback is not taken into account in the FN approach. Similarly, DOC release from phytoplankton and heterotrophic bacteria is systematically higher in the FN models. While the mechanism generating this difference is less obvious, it likely reflects the dynamic adjustment of biomass and fluxes in the DS approach, in contrast to the fixed flux partitioning used in the FN models.

**Figure 5.**
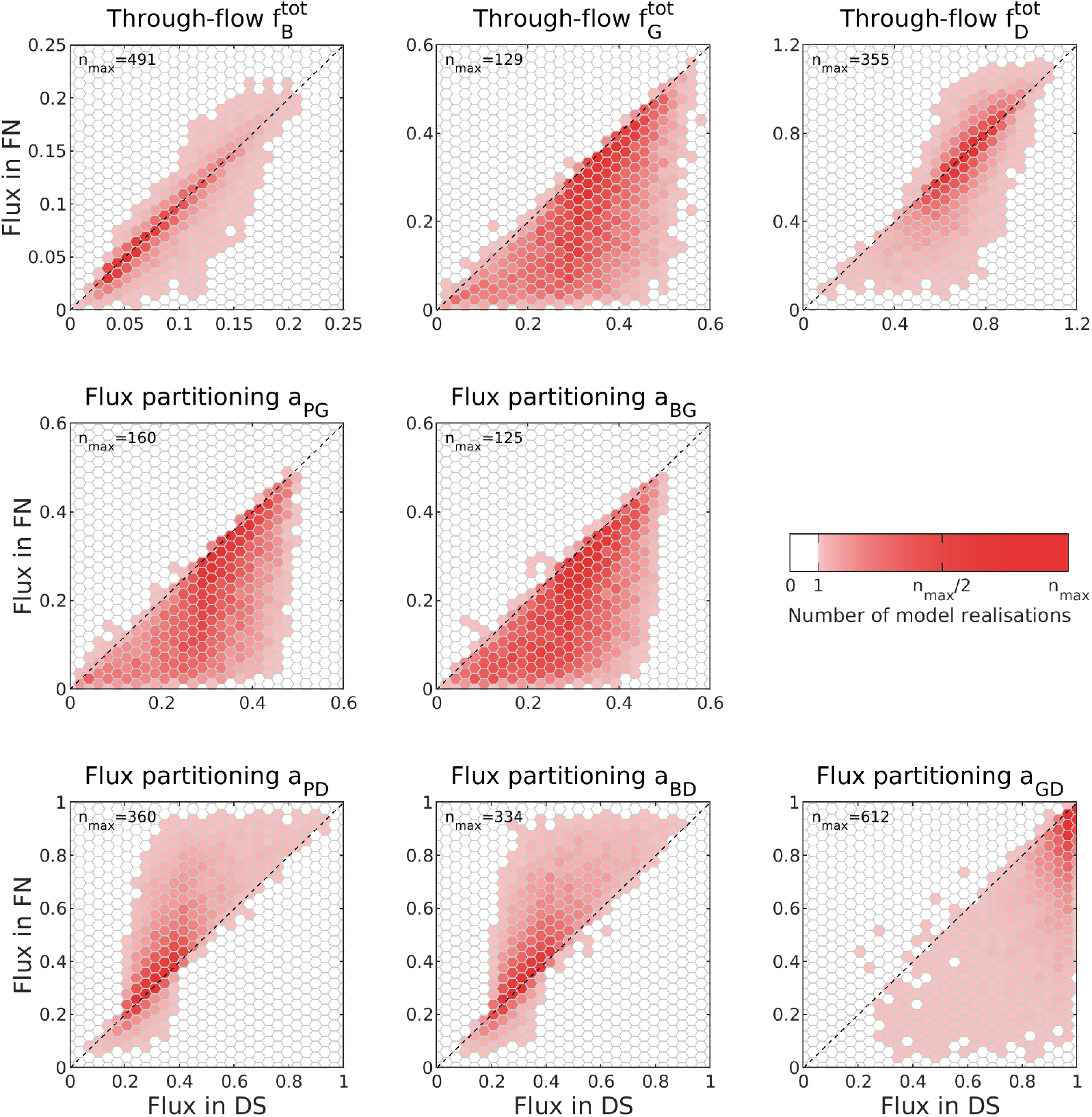
Comparison of through-flow fluxes and partitioning ratios between the DS and FN models following virus removal. In all panels, DS model fluxes are shown on the horizontal axes, and FN model fluxes on the vertical axes. The top row shows through-flow fluxes for heterotrophic bacteria, grazers and DOC (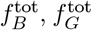 and 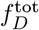, respectively). The middle row shows grazing partitioning ratios for phytoplankton (left) and heterotrophic bacteria (right). The bottom row shows partitioning ratios of DOC release from phytoplankton, heterotrophic bacteria and grazers. Colors indicate the number of model realisations in each hexagonal bin, normalised by the maximum bin count *n*_max_, which is indicated in each panel.

Overall, these results indicate that while the FN model reproduces some aspects of the system response, it neglects several feedback mechanisms that are present in the DS model and that can strongly influence the outcomes of virus removal. In particular, the fixed partitioning ratios and the absence of compartment stock dynamics in the FN approach limit its ability to represent the dynamical responses and indirect interactions of the DS approach. In contrast, DS models enable a more comprehensive understanding of the full range of system-level effects of virus removal.

## 6 Discussion

In this paper we connected existing quantitative approaches to assess the impact of viral lysis on marine carbon fluxes within a common theoretical framework. Dynamical system (DS) models (Weitz *et al*., 2015), flux network (FN) models (Wilhelm & Suttle, 1999), and on-board experimental approaches such as modified dilution (MD) and virus reduction (VR) assays (Mojica, 2015), address virus–microbe–biogeochemistry coupling at different scales, with distinct strengths and limitations. As a result, they provide non-equivalent but complementary quantifications of marine viral lysis and its role in ocean carbon cycling. To illustrate this, we performed different analyses and first showed that commonly used lysis percentages refer to fundamentally different quantities: short-term process-based rates versus system-level flux ratios, and cannot be directly compared. These measures can diverge strongly depending on the system productivity and recycling intensity while having identical lysis strengths. We further compared published DS and FN models that include viral lysis. We showed a large variability in predicted viral impacts that is driven primarily by parameterisation rather than by modelling framework. Finally, we evaluated the modelling practice of virus removal as a diagnostic tool of the impact of viruses on a system. We showed that lysis fluxes poorly predict the impacts of virus removal, as carbon pathways reorganise in response to virus removal. Additionally, dynamical feedbacks captured in DS models but absent from FN models can strongly influence the outcomes of removing viruses.

Placing our results in the context of reported ranges highlights the large variability in estimated viral impacts across both in situ measurements and model-based approaches. For process-based lysis percentages, Mojica & Brussaard (2026) reported phytoplankton viral lysis ranging from 0.7 to 25% d^−1^ (25th − 75th percentiles; see Supporting Information Fig. S4, MD assays) and heterotrophic bacterial lysis rates ranging from 35 to 140% d^−1^ (25th − 75th percentiles; VR assays). For flux-based lysis percentages, our model meta-analysis yielded lysis flux ratios ranging from 12 to 80% of primary production, indicating greater variability than the still widely cited range of 6 to 26% of primary production reported by Wilhelm & Suttle (1999). Beyond differences between metrics, the large parameter dispersion revealed by our PCA analysis of DS and FN model outputs indicates substantial variability across models. Rather than reflecting only methodological uncertainty, this variability likely captures real ecological heterogeneity across systems, including differences between oligotrophic and coastal systems, seasonal succession and depth gradients. Dissecting variability along these environmental and trophic gradients offers a promising path toward reconciling divergent estimates and identifying regime-dependent controls on viral impacts. In this context, the synthesis by Mojica & Brussaard (2026) represents an important step toward quantifying viral lysis across ecosystem types, and highlights the need for closer integration between observational datasets and ecosystem-scale models to constrain viral process rates across environmental regimes.

The quantitative approaches tested here also differ fundamentally in the temporal and organisational scales they resolve, and thus in the types of viral impacts they can capture. Experimental on-board methods provide direct, empirical quantification of virus-mediated mortality and associated carbon fluxes at local scales. They resolve short-term virus–host dynamics, which can vary over diel (Aylward *et al*., 2017, Beckett *et al*., 2024) to seasonal timescales (Biggs *et al*., 2021, Bolaños *et al*., 2024). New molecular in situ methods of viral lysis or infectivity such as MORS (Zhong *et al*., 2023), VirusFISH (Castillo *et al*., 2020) or iPolony (Mruwat *et al*., 2021) can provide complementary and more direct information on virus–host interactions, but they are still limited to local and short-time scales. Additionally, these methods capture only a subset of ecosystem processes and should be integrated with measurements or modelling of non-viral pathways to place viral impacts in a system-wide context (Mateus, 2017, Talmy *et al*., 2019, Beckett *et al*., 2024). Flux network models offer a steady-state representation of carbon partitioning across the microbial system, and can be used to connect short-term flux partitioning ratios inferred from experiments to long-term equilibrium fluxes (Xie *et al*., 2022). They are generally simpler to implement than dynamical models and still provide a system-level quantification of how viral lysis and grazing redistribute carbon, but they lack explicit dynamics, feedbacks, stock concentrations and turnover times. In contrast, dynamical system models explicitly represent population and carbon stock dynamics through time, allowing mechanistic exploration on how microbial host mortality is partitioned between viral lysis and grazing and how it changes with environmental drivers across multiple temporal and spatial scales. This enables analyses of perturbations such as heat waves (Soulié *et al*., 2022, 2023) or nutrient limitation (Maat & Brussaard, 2016, Flynn *et al*., 2021, Yung *et al*., 2025) and of system responses to changes in viral pressure (Berdjeb *et al*., 2011). However, these advantages come at the cost of substantial parameter uncertainty and reliance on strong structural assumptions or experimental calibration.

Finally, connecting these approaches can help in reducing the uncertainties but requires explicit integration across scales to inform system-level carbon budgets and long-term ecosystem functioning. As emphasized by Vincent & Vardi (2023), a comprehensive assessment of the viral shunt requires a multi-scale perspective that links processes measured at the bottle or station scale to emergent, large-scale carbon fluxes. Additionally, viral impacts on carbon cycling extend beyond the viral shunt to include effects on particle formation, aggregation and export, commonly referred as the viral shuttle (Weinbauer, 2004, Yamada *et al*., 2018, Kaneko *et al*., 2021). These apparent opposing pathways imply that the net effect of viruses on carbon export depends critically on how viral processes are represented within ecosystem and biogeochemical frameworks. New efforts to incorporate viral processes into global-scale models explore new ways of representing viral effects, including metabolic modelling approaches that link viral activity to metabolite fluxes (Régimbeau *et al*., 2025, Talmy *et al*., 2025). In contrast, the approach adopted here focuses on reconciling classical process-based and flux-based representations of viral lysis across local to ecosystem scales, with the goal of clarifying how different methodological choices shape inferred viral impacts. These perspectives are complementary: large-scale models require empirically and mechanistically grounded constraints on viral flux partitioning, while ecosystem-level and experimental studies benefit from embedding their results within global modelling frameworks (Frémont *et al*., 2025). Progress toward a predictive representation of viral impacts on marine carbon cycling will therefore require tighter coupling between quantitative experimental methods, ecosystem-level and global-scale biogeochemical models.

## Acknowledgments

MP acknowledges financial support from the Ocean Institute of Sorbonne University Alliance through a PhD grant (Grant No. U23JR31001-1).

