## Supporting Information for "Connecting models, networks and experiments: Revisiting the role of viruses in marine carbon cycling"

### S1 A minimal microbial ecosystem model

Here we introduce the microbial ecosystem model that is referred to in the main text. It is a simplified version of the model studied by Weitz *et al.* (2015). The main differences are:

- Whereas Weitz *et al.* (2015) distinguish cyanobacteria and eukaryotic phytoplankton, here we aggregate all primary producers into one dynamical variable $P$. Similarly, we do not differentiate viruses infecting cyanobacteria from those infecting eukaryotic phytoplankton, but instead consider a single virus population $V_P$ infecting all phytoplankton.
- Weitz *et al.* (2015) focus on nitrogen cycling within the microbial ecosystem, resulting in a model that is effectively closed with respect to this nutrient. Their model includes two abiotic compartments, organic and inorganic nutrients, that limit different populations. In contrast, we focus on carbon flows. Our model includes only one abiotic compartment: dissolved organic carbon. We do not track inorganic nutrients and assume that inorganic carbon is never limiting.
- In the model of Weitz *et al.* (2015) primary production is limited by the availability of inorganic nitrogen. Here we replace this limitation with self-limitation, i.e., we assume logistic growth for the phytoplankton population with a fixed carrying capacity $K$. Consequently, we do not account for the positive effect of remineralisation on phytoplankton growth.

As a result, our minimal microbial ecosystem model comprises six dynamical variables (see Table S1): phytoplankton $P$, heterotrophic bacteria $B$, zooplankton grazers $G$ (consuming both phytoplankton and heterotrophic bacteria), phytoplankton viruses $V_P$, bacteriophages $V_B$, and dissolved organic carbon $D$. Biotic compartments are expressed as abundances (number of individuals per volume unit). Because we assume fixed carbon quotas for all compartments, these variables can equivalently be expressed in carbon units.

The dynamical equations are obtained by enumerating the processes that affect each compartment. Using the parameters listed in Table S2, we get

$$\frac{\mathrm{d}P}{\mathrm{d}t} = \mu_P(P)P - \psi_{PG}\,PG - \phi_P P V_P - \ell_P^{\mathrm{inorg}} P - \ell_P^{\mathrm{org}} P \tag{S1a}$$

$$\frac{\mathrm{d}B}{\mathrm{d}t} = \mu_B(D)B - \psi_{BG}\,BG - \phi_B B V_B - \ell_B^{\mathrm{inorg}} B - \ell_B^{\mathrm{org}} B \tag{S1b}$$

$$\frac{\mathrm{d}G}{\mathrm{d}t} = p_G^{\mathrm{assim}}\psi_{PG}\, PG\frac{q_P}{q_G} + p_G^{\mathrm{assim}}\psi_{BG}\, BG\frac{q_B}{q_G} - \ell_G^{\mathrm{inorg}}G - \ell_G^{\mathrm{cons}}G^2 \tag{S1c}$$

$$\frac{\mathrm{d}V_P}{\mathrm{d}t} = \beta_P\phi_P PV_P - \ell_{V_P}V_P \tag{S1d}$$

$$\frac{\mathrm{d}V_B}{\mathrm{d}t} = \beta_B\phi_B BV_B - \ell_{V_B}V_B \tag{S1e}$$

$$\begin{aligned}\frac{\mathrm{d}D}{\mathrm{d}t} = {} & p_G^{\mathrm{org}}\,\psi_{PG}\, PG\, q_P + \ell_P^{\mathrm{org}}P\, q_P + p_G^{\mathrm{org}}\,\psi_{BG}\, BG\, q_B + \ell_B^{\mathrm{org}}B\, q_B \\ & + \ell_G^{\mathrm{cons}}G^2\, q_G + \ell_{V_P}V_P\, q_V + \ell_{V_B}V_B\, q_V \\ & + \phi_P PV_P\,(q_P - \beta_P q_V) + \phi_B BV_B\,(q_B - \beta_B q_V) \\ & - \frac{1}{p_{DB}^{\mathrm{assim}}}\mu_B(D)B\, q_B\end{aligned} \tag{S1f}$$

The growth functions for phytoplankton and for heterotrophic bacteria are

$$\mu_P(P) = \mu_P^{\max}\Big(1 - \frac{P}{K_P}\Big) \qquad \text{and} \qquad \mu_B(D) = \mu_B^{\max}\frac{B}{K_{DB} + D}$$

We focus on the equilibrium state of the model, which can be determined analytically. Because viruses exert top-down control on their hosts, the equilibrium values of $P$ and $B$ follow directly from the conditions $\frac{\mathrm{d}V_P}{\mathrm{d}t} = 0$ and $\frac{\mathrm{d}V_B}{\mathrm{d}t} = 0$. These values then allow us to determine $G$ from $\frac{\mathrm{d}G}{\mathrm{d}t} = 0$ and $V_P$ from $\frac{\mathrm{d}P}{\mathrm{d}t} = 0$. Finally, the remaining variables, $V_B$ and $D$, can be obtained from $\frac{\mathrm{d}B}{\mathrm{d}t} = 0$ and $\frac{\mathrm{d}D}{\mathrm{d}t} = 0$. This gives the following equilibrium expressions:

$$P = \frac{\ell_{V_P}}{\beta_P\phi_P} \tag{S2a}$$

$$B = \frac{\ell_{V_B}}{\beta_B\phi_B} \tag{S2b}$$

$$G = \frac{1}{\ell_G^{\mathrm{cons}}}\Bigg(p_G^{\mathrm{org}}\Big(\frac{q_P}{q_G}\psi_{PG}P + \frac{q_B}{q_G}\psi_{BG}B\Big) - \ell_G^{\mathrm{inorg}}\Bigg) \tag{S2c}$$

$$V_P = \frac{1}{\phi_P}\Bigg(\mu_P^{\max}\Big(1 - \frac{P}{K_P}\Big) - \psi_{PG}G - \ell_P\Bigg) \tag{S2d}$$

$$V_B = \frac{\psi_{BG}G + \ell_B - p_{DB}^{\mathrm{assim}}x_1}{1 - p_{DB}^{\mathrm{assim}}} \tag{S2e}$$

$$D = \frac{x_2}{\mu_B^{\max} - x_2}K_{DB} \tag{S2f}$$

where we used the following short-hand notation:

$$\begin{aligned} x_1 &= c_{\frac{P}{B}}\Big(\phi_P V_P + p_G^{\mathrm{org}}\psi_{PG}G + \ell_P^{\mathrm{org}}\Big) + p_G^{\mathrm{org}}\psi_{BG}G + \ell_B^{\mathrm{org}} + c_{\frac{G}{B}}\ell_G^{\mathrm{cons}}G \\ x_2 &= \phi_B V_B + \psi_{BG}G + \ell_B \\ \ell_P &= \ell_P^{\mathrm{inorg}} + \ell_P^{\mathrm{org}} \\ \ell_B &= \ell_B^{\mathrm{inorg}} + \ell_B^{\mathrm{org}} \\ c_{\frac{P}{B}} &= \frac{q_P P}{q_B B} \end{aligned}$$

Table S1: Variables of model (S1) with units and value ranges.

| symbol | meaning | units | lower bound | upper bound |
|---|---|---|---|---|
| $P$ | phytoplankton abundance | cells $L^{-1}$ | $2 \times 10^5$ | $2 \times 10^7$ |
| $B$ | abundance of heterotrophic bacteria | cells $L^{-1}$ | $2 \times 10^7$ | $2 \times 10^9$ |
| $G$ | abundance of zooplankton grazers | individuals $L^{-1}$ | $2 \times 10^3$ | $2 \times 10^5$ |
| $V_P$ | abundance of phytoplankton viruses | viruses $L^{-1}$ | $2 \times 10^6$ | $2 \times 10^8$ |
| $V_B$ | abundance of bacteriophages | viruses $L^{-1}$ | $2 \times 10^8$ | $2 \times 10^{10}$ |
| $D$ | dissolved organic carbon concentration | $\mu$mol-C $L^{-1}$ | 30 | 100 |

$$c_{\frac{G}{B}} = \frac{q_G G}{q_B B}$$

To generate realisations of the model, we first sample parameter values. Table S2 lists for each parameter a range of realistic values, that we adapted from those reported in Weitz *et al.* (2015). Parameter values are sampled on a logarithmic scale; that is, we draw the logarithm of each parameter value uniformly from the range defined by the logarithms of its lower and upper bounds. For a given set of parameter values, we evaluate the equilibrium using eqs. (S2). For each of the six dynamical variables, we verify whether the equilibrium value lies in between the bounds given in Table S1, which were also adapted from those reported in Weitz *et al.* (2015). Realisations satisfying this condition for all six variables are accepted; otherwise, they are rejected.

### S2 Analysis of flux network models

The flux network approach enables the reconstruction of stationary carbon fluxes in the microbial ecosystem using only a limited amount of quantitative information. We denote by $F_{ij}$ the flux from compartment $i$ to compartment $j$, and by $F_i^{\text{tot}}$ the through-flow of compartment $i$, defined as the sum of all incoming fluxes, which, due to stationarity, is equal to the sum of all outgoing fluxes. The approach assumes that for each compartment the through-flow is distributed in fixed proportions among the different outgoing links. This distribution is specified by partitioning ratios $a_{ij}$, defined as:

$$a_{ij} = \frac{F_{ij}}{F_i^{\text{tot}}}, \tag{S3}$$

For illustration, consider the minimal microbial ecosystem model introduced in Sect. S1. We label the compartments as follows:

| | |
|---|---|
| $i = 1$ | phytoplankton $P$ |
| $i = 2$ | heterotrophic bacteria $B$ |
| $i = 3$ | zooplankton grazers $G$ |
| $i = 4$ | dissolved organic carbon $D$ |

Primary production, i.e., the through-flow $F_1^{\text{tot}} = F_P^{\text{tot}}$, is directed toward the grazer and DOC compartments. The corresponding flux partitioning could be:

$$a_{13} = a_{PG} = 0.50 \qquad a_{14} = a_{PD} = 0.40 \qquad a_{1j} = 0 \text{ for other compartments } j$$

Table S2: Parameters of model (S1) with units and value ranges.

| symbol | meaning | units | lower bound | upper bound |
|---|---|---|---|---|
| $q_P$ | carbon quota of $P$ | $\mu$mol-C/cell | $3 \times 10^{-7}$ | $3 \times 10^{-6}$ |
| $q_B$ | carbon quota of $B$ | $\mu$mol-C/cell | $2 \times 10^{-9}$ | $2 \times 10^{-8}$ |
| $q_G$ | carbon quota of $G$ | $\mu$mol-C/ind | $2 \times 10^{-4}$ | $2 \times 10^{-3}$ |
| $q_V$ | carbon quota of $V_P$ and $V_B$[a] | $\mu$mol-C/virus | $2 \times 10^{-12}$ | $6 \times 10^{-11}$ |
| $\mu_P^{\max}$ | maximal growth rate of $P$ | $d^{-1}$ | 0.2 | 2.0 |
| $K_P$ | carrying capacity of $P$ | cell $L^{-1}$ | $1 \times 10^{7}$ | $1 \times 10^{9}$ |
| $\ell_P^{\text{inorg}}$ | loss rate of $P$ as inorganic carbon | $d^{-1}$ | $1 \times 10^{-3}$ | 0.1 |
| $\ell_P^{\text{org}}$ | loss rate of $P$ as organic carbon | $d^{-1}$ | $5 \times 10^{-3}$ | 0.1 |
| $\psi_{PG}$ | grazing rate of $P$ by $G$ | L/ind $d^{-1}$ | $1 \times 10^{-6}$ | $2 \times 10^{-5}$ |
| $\mu_B^{\max}$ | maximal growth rate of $B$ | $d^{-1}$ | 0.5 | 2.0 |
| $K_{BD}$ | half-saturation constant of $B$ consuming $D$ | $\mu$mol-C $L^{-1}$ | 30 | 300 |
| $p_{BD}^{\text{assim}}$ | assimilation efficiency[b] of $B$ consuming $D$ | — | 0.05 | 0.2 |
| $\ell_B^{\text{inorg}}$ | loss rate of $B$ as inorganic carbon | $d^{-1}$ | $1 \times 10^{-3}$ | 0.1 |
| $\ell_B^{\text{org}}$ | loss rate of $B$ as organic carbon | $d^{-1}$ | $5 \times 10^{-3}$ | 0.1 |
| $\psi_{BG}$ | grazing rate of $B$ by $G$ | L/ind $d^{-1}$ | $1 \times 10^{-6}$ | $3 \times 10^{-5}$ |
| $\ell_G^{\text{inorg}}$ | loss rate of $G$ as inorganic carbon | $d^{-1}$ | $1 \times 10^{-3}$ | $1 \times 10^{-2}$ |
| $\ell_G^{\text{cons}}$ | coefficient of quadratic loss term[c] of $G$ | L/ind $d^{-1}$ | $1 \times 10^{-7}$ | $1 \times 10^{-5}$ |
| $p_G^{\text{assim}}$ | assimilation efficiency[b] of $G$ | — | 0.3 | 0.5 |
| $p_G^{\text{org}}$ | fraction of grazing lost as organic carbon[d] | — | 0.2 | 0.4 |
| $\phi_P$ | infection rate of $V_P$ | L/virus $d^{-1}$ | $10^{-9}$ | $10^{-8}$ |
| $\beta_P$ | burst size of $V_P$ | — | 50 | 500 |
| $\ell_{V_P}$ | loss rate of $V_P$ | $d^{-1}$ | 0.05 | 5.0 |
| $\phi_B$ | infection rate of $V_B$ | L/virus $d^{-1}$ | $10^{-11}$ | $10^{-9}$ |
| $\beta_B$ | burst size of $V_B$ | — | 10 | 100 |
| $\ell_{V_B}$ | loss rate of $V_B$ | $d^{-1}$ | 0.05 | 5.0 |

[a] The carbon quotas of phytoplankton viruses and bacteriophages are assumed to be equal.

[b] The assimilation efficiency of a consumption process (i.e., $D$ by $B$, $P$ by $G$ and $B$ by $G$) is defined as the fraction of carbon assimilated relative to carbon consumed. The assimilation efficiencies of $P$ by $G$ and of $B$ by $G$ are assumed to be equal.

[c] Generally, a quadratic loss term serves as an approximation of the consumption of $G$ by higher trophic levels. Here, for simplicity, we consider only the fraction of the consumed carbon that ultimately returns to the dissolved organic carbon pool.

[d] Fraction of carbon consumed by grazers that is lost as dissolved organic carbon. The loss fractions of $G$ consuming $P$ and $G$ consuming $B$ are assumed to be equal.

The remaining fraction $1 - \sum_j a_{1j} = 0.10$ represents respiratory losses (carbon leaving the system as $CO_2$). The partitioning ratios satisfy the inequality $\sum_j a_{ij} \leq 1$, because the total flux leaving a compartment cannot be larger than the flux entering it. We denote the loss flux by

$$F_{i0} = F_i^{\text{tot}} - \sum_j F_{ij} = F_i^{\text{tot}} \Big(1 - \sum_j a_{ij}\Big),$$

where the subscipt 0 denotes the external environment.

The partitioning ratios $a_{ij}$ define the structure of the flux network and determine the distribution of carbon fluxes within it. To reconstruct the network, it suffices to compute the through-flows $F_i^{\text{tot}}$, because once these are known, eq. (S3) can be used to compute the fluxes $F_{ij}$ between compartments. The through-flows satisfy the following balance equation:

$$F_j^{\text{tot}} = \sum_i F_i^{\text{tot}} a_{ij} \tag{S4}$$

This equation states that the total through-flow of compartment $j$ (left-hand side) is equal to the sum of all fluxes entering that compartment (right-hand side).

The balance equation (S4) applies to all compartments except $i = 1$. This exception arises because the equation does not account for carbon fixation, by which carbon is taken up directly from the environment. We denote this input flux by $F_{01}$, where the subscript 0 again refers to the environment. Hence, we obtain the following system of equations for the through-flows $F_i^{\text{tot}}$:

$$\begin{aligned} F_1^{\text{tot}} &= F_{01} + \sum_i F_i^{\text{tot}} a_{i1} \\ F_j^{\text{tot}} &= \sum_i F_i^{\text{tot}} a_{ij} \qquad \text{for } j \geq 2, \end{aligned}$$

or in matrix form,

$$\underbrace{\vec{v}_F}_{\substack{\text{row}\\\text{vector}}} \underbrace{(\mathbb{1} - A)}_{\text{matrix}} = F_{01}\, \vec{e}_1 \tag{S5}$$

with $\vec{v}_F$ the vector of through-flows, $A$ the partitioning ratio matrix, $\mathbb{1}$ the identity matrix, and $\vec{e}_1$ the first unit vector (first component equal to one; all other components equal to zero),

$$\vec{v}_F = \begin{pmatrix} F_1^{\text{tot}} & F_2^{\text{tot}} & F_3^{\text{tot}} & \cdots \end{pmatrix} \qquad A = \begin{pmatrix} 0 & a_{12} & a_{13} & \cdots \\ a_{21} & 0 & a_{23} & \cdots \\ a_{31} & a_{32} & 0 & \cdots \\ \vdots & \vdots & \vdots & \end{pmatrix}$$

Here the primary production $F_{01}$ serves as a scaling factor. Typically, primary production is used as the reference for carbon flux, so that all fluxes $F_i^{\text{tot}}$ and $F_{ij}$ within the system are expressed relative to it.

The solution of eq. (S5) is given by

$$\vec{v}_F = F_{01}\, \vec{e}_1\, (\mathbb{1} - A)^{-1} = F_{01} \Big(\vec{e}_1 + \vec{e}_1 A + \vec{e}_1 A^2 + \vec{e}_1 A^3 + \ldots\Big) \tag{S6}$$

The latter expression shows that the through-flows $F_i^{\text{tot}}$ result from the sum of a large number of flux pathways originating from primary production. In particular, the same quantity of fixed carbon can contribute multiple times to a given through-flow. Hence, a through-flow should not be interpreted as a proportion of primary production, but rather as a flux expressed in units of primary production. The same interpretation applies to the fluxes $F_{ij}$ between compartments.

Finally, we note that the equilibrium of the minimal microbial ecosystem model of Sect. S1 can be represented as a flux network model. This flux network model has four compartments: $P$, $B$, $G$ and $D$. The inter-compartment fluxes $F_{ij}$ and through-flow fluxes $F_i^{\text{tot}}$ are given by

$$
\begin{aligned}
F_P^{\text{tot}} &= \mu_P(P)P\, q_P \\
F_{PG} &= p_G^{\text{assim}}\, \psi_{PG}\, PG\, q_P \\
F_{PD} &= F_{PD}^0 + F_{PD}^V \\
F_{PD}^0 &= p_G^{\text{org}}\, \psi_{PG}\, PG\, q_P + \ell_P^{\text{org}} P\, q_P \\
F_{PD}^V &= \ell_{V_P} V_P\, q_V + \phi_P P V_P\, (q_P - \beta_P q_V) = \phi_P P V_P q_P \\
F_B^{\text{tot}} &= \mu_B(D)B\, q_B \\
F_{BG} &= p_G^{\text{assim}} \psi_{BG}\, BG\, q_B \\
F_{BD} &= F_{BD}^0 + F_{BD}^V \\
F_{BD}^0 &= p_G^{\text{org}}\, \psi_{BG}\, BG\, q_B + \ell_B^{\text{org}} B\, q_B \\
F_{BD}^V &= \ell_{V_B} V_B\, q_V + \phi_B B V_B\, (q_B - \beta_B q_V) = \phi_B B V_B q_B \\
F_G^{\text{tot}} &= p_G^{\text{assim}} \psi_{PG}\, PG\, q_P + p_G^{\text{assim}} \psi_{BG}\, BG\, q_B \\
F_{GD} &= \ell_G^{\text{cons}} G^2\, q_G \\
F_D^{\text{tot}} &= \frac{1}{p_{DB}^{\text{assim}}} \mu_B(D)B\, q_B \\
F_{DB} &= \mu_B(D)B\, q_B
\end{aligned}
$$

Here the fluxes $F_{PD}$ and $F_{BD}$ are decomposed into two components: a virus-independent contribution ($F_{PD}^0$ and $F_{BD}^0$) and a virus-mediated contribution ($F_{PD}^V$ and $F_{BD}^V$).

### S3 Quantitative analysis of MD/VR assays

Here we briefly present a quantitative treatment of the modified dilution and virus reduction assays. We adopt a principled perspective: we assume that the various potential biases can be neglected, in order to focus on how the measured quantities relate to microbial ecosystem models.

#### Modified dilution assay

The modified dilution (MD) assay is based on measuring the phytoplankton growth rate in grazer-free (in which grazers are removed) and virus-free treatments (in which both grazers and viruses are removed). To connect this setup to microbial ecosystem models, we consider the dynamical equation for phytoplankton $P$ of our minimal model, eqs. (S1a),

$$
\frac{\mathrm{d}P}{\mathrm{d}t} = \mu_P(P)P - \psi_{PG}PG - \phi_P P V_P - \ell_P P
$$

$$= \Big(\mu_P(P) - \psi_{PG}G - \phi_P V_P - \ell_P\Big)P$$

Recall that the dependence of the gross growth rate $\mu_P(P)$ on population size $P$ corresponds to density-dependent growth.

The net growth rate of phytoplankton is measured in two dilution series experiments,

(1) $P$ and $G$ are diluted by a dilution factor $\gamma$, while $V_P$ is kept unchanged. Hence, we find for the net growth rate,

$$g_P^{(1)} = \mu_P(\gamma P) - \psi_{PG}\gamma G - \phi_P V_P - \ell_P$$

(2) $P$, $G$ and $V_P$ are diluted by a factor $\gamma$. Hence, the net growth rate is

$$g_P^{(2)} = \mu_P(\gamma P) - \psi_{PG}\gamma G - \phi_P \gamma V_P - \ell_P$$

We assume that $\mu_P(P)$ depends only weakly on $P$, so that the dependence of $\mu_P(\gamma P)$ on $\gamma$ can be neglected. Then,

$$\begin{aligned} &\text{the regression slope of } g_P^{(1)} \text{ vs } \gamma \text{ gives an estimate of } \psi_{PG}G \\ &\text{the regression slope of } g_P^{(2)} \text{ vs } \gamma \text{ gives an estimate of } \psi_{PG}G + \phi_P V_P \end{aligned}$$

Subtracting the first from the second estimate gives an estimate of $\phi_P V_P$. That is, the assay yields estimates of the grazing and virus-induced mortality rates of phytoplankton. Combining this with estimates of the phytoplankton abundance $P$ and carbon quota $q_P$, we get estimates of the fluxes

$$F_{PG} = \psi_P PG\, q_P \qquad \text{and} \qquad F_{PD}^V = \phi_P P V_P\, q_P$$

In practice, the conditions of the MD assay do not match the idealised assumptions used above:

- The reduction in phytoplankton density during dilution can alter density-dependent growth processes, leading to biased estimates.
- The simultaneous dilution of dissolved organic carbon and inorganic nutrients can modify resource availability and thus influence the microbial dynamics.
- Natural communities contain a diversity of hosts and virus types. Representing this diversity with a single effective host–virus pair may be inaccurate (Beckett & Weitz, 2018).

### Virus reduction assay

To analyse the quantitative details of the virus reduction (VR) assay, we extend the virus-host dynamics of heterotrophic bacteria by distinguishing between uninfected and infected bacteria. We write $B = B_U + B_I$ where $B_U$ and $B_I$ denote the uninfected (susceptible) and infected bacterial populations, respectively. After excluding grazers, their dynamics are

$$\frac{\mathrm{d}B_U}{\mathrm{d}t} = \mu_B(D)B_U - \phi_B V_B B_U - \ell_B B_U$$
$$\frac{\mathrm{d}B_I}{\mathrm{d}t} = \phi_B V_B B_U - \eta_B B_I - \ell_B B_I$$

$$\frac{\mathrm{d}V_B}{\mathrm{d}t} = \beta_B \eta_B B_I - \phi_B V_B B_U - \ell_{V_B} V_B$$

where $\phi_B$ is the infection rate, $\eta_B$ the lysis rate (the reciprocal of latent period) and $\beta_B$ the burst size.

Ideally, the virus reduction step removes all free viruses from the system while retaining the viruses associated with infected hosts. The dynamics for the incubation of viruses and hosts are

$$\begin{aligned}
\frac{\mathrm{d}B_U}{\mathrm{d}t} &= \mu_B(D) B_U - \ell_B B_U \\
\frac{\mathrm{d}B_I}{\mathrm{d}t} &= -\eta_B B_I - \ell_B B_I \\
\frac{\mathrm{d}V_B}{\mathrm{d}t} &= \beta_B \eta_B B_I
\end{aligned}$$

We see that the initial increase in virus abundance is due to already infected cells finishing their replication cycle and lysing. The slope of this increase provides an estimate of the product $\beta_B \eta_B B_I$. Given an independent estimate of the burst size $\beta_B$, we get an estimate for $\eta_B B_I$, which represents the virus-induced loss rate of the bacteria compartment,

$$\frac{\mathrm{d}(B_U + B_I)}{\mathrm{d}t} = \mu_B(D) B_U - \underbrace{\eta_B B_I}_{\substack{\text{virus-induced} \\ \text{mortality}}} - \ell_B (B_U + B_I)$$

Relating this back to the dynamics of bacteria and their viruses in the original model, eq. (S1b),

$$\frac{\mathrm{d}B}{\mathrm{d}t} = \mu_B(D) B - \underbrace{\phi_B B V_B}_{\substack{\text{virus-induced} \\ \text{mortality}}} - \ell_B B$$

we see that the estimated virus-induced loss rate corresponds to the product $\phi_B B V_B$. Multiplying by the bacteria carbon quota $q_B$, we obtain an estimate of the flux

$$F^V_{BD} = \phi_B B V_B \, q_B$$

In practice, departures from the idealised assumptions can affect the estimates:

- Uncertainties in the burst size directly propagate to the estimated flux, since this parameter enters linearly into the calculation.
- Natural microbial communities contain multiple host–virus pairs. Assuming a single representative burst size may be inaccurate.
- Non-lytic, temperate or chronic infections are not represented in the idealised model.
- Secondary infections may occur during incubation if its duration approaches the latent period.
- The exclusion of grazers can modify the bacteria dynamics.

### S4 Lysis fluxes and carbon recycling

In Section 3 we presented an example illustrating the potential interaction between carbon recycling within the microbial food web and virus-induced lysis fluxes. Here we provide the full details of this example.

We introduce a partitioning ratio matrix in which virus-mediated fluxes are isolated from all other fluxes. Starting from the minimal microbial ecosystem of Sect. S1, we add a (virtual) compartment $D^V$ through which the lysis fluxes pass before being transferred to the DOC compartment. Ordering the compartments as $P, B, G, D, D^V$, the partitioning ratio matrix is given by

$$A = \begin{pmatrix} 0 & 0 & a_{PG} & a^0_{PD} & a^V_{PD} \\ 0 & 0 & a_{BG} & a^0_{BD} & a^V_{BD} \\ 0 & 0 & 0 & a_{GD} & 0 \\ 0 & a_{DB} & 0 & 0 & 0 \\ 0 & 0 & 0 & 1 & 0 \end{pmatrix} \tag{S7}$$

Here the partitioning ratios $a_{PD}$ and $a_{BD}$ are decomposed into two components,

$$a_{PD} = a^0_{PD} + a^V_{PD} \qquad \text{and} \qquad a_{BD} = a^0_{BD} + a^V_{BD}$$

where $a^0_{PD}$ and $a^0_{BD}$ represent virus-independent contributions, and $a^V_{PD}$ and $a^V_{BD}$ represent virus-mediated contributions. Note in eq. (S7) the entry equal to 1 in the last row of the matrix $A$, which corresponds to the transfer of the lysis-derived fluxes to the DOC compartment.

To vary carbon cycling within the ecosystem, we keep all partitioning ratios constant except for $a_{DB}$, which can be interpreted as the efficiency with which bacteria consume DOC. Specifically, we set

$$A = \begin{pmatrix} 0 & 0 & 0.30 & 0.30 & 0.30 \\ 0 & 0 & 0.20 & 0.10 & 0.70 \\ 0 & 0 & 0 & 0.20 & 0 \\ 0 & a_{DB} & 0 & 0 & 0 \\ 0 & 0 & 0 & 1 & 0 \end{pmatrix}$$

and examine different values for $a_{DB}$:

- $a_{DB} = 0$ or no recycling (theoretical case): All DOC through-flow is lost from the microbial loop. The stationary fluxes correspond to fixed carbon passing once through the compartments:

  $$f^{\text{tot}}_D = 0.66 \qquad f^{\text{tot}}_B = 0 \qquad f^V_{PD} = 0.30 \qquad f^V_{BD} = 0$$

  where the fluxes have been normalised by primary production,

  $$f^{\text{tot}}_i = \frac{F^{\text{tot}}_i}{F^{\text{tot}}_P} \qquad \text{and} \qquad f_{ij} = \frac{F_{ij}}{F^{\text{tot}}_P}.$$

- $a_{DB} = 0.10$ or weak recycling: Only 10% of the DOC through-flow is retained in the microbial loop, so recycling is minimal and stationary fluxes remain close to the first-passage values:

  $$f^{\text{tot}}_D = 0.72 \qquad f^{\text{tot}}_B = 0.07 \qquad f^V_{PD} = 0.30 \qquad f^V_{BD} = 0.05$$

- $a_{DB} = 0.40$ or strong recycling: With 40% recycling efficiency, carbon circulates several times before exiting, increasing stationary fluxes:

$$f_D^{\text{tot}} = 0.99 \qquad f_B^{\text{tot}} = 0.40 \qquad f_{PD}^V = 0.30 \qquad f_{BD}^V = 0.28$$

- $a_{DB} = 0.70$ or very strong recycling: Only 30% of the DOC through-flow is lost from the microbial loop, producing large stationary fluxes dominated by recycled carbon rather than first-passage contributions:

$$f_D^{\text{tot}} = 1.60 \qquad f_B^{\text{tot}} = 1.12 \qquad f_{PD}^V = 0.30 \qquad f_{BD}^V = 0.78$$

- $a_{DB} = 1$, or perfect recycling (theoretical case): The entire DOC through-flow is consumed by bacteria and returned to the microbial loop, yielding maximal stationary fluxes:

$$f_D^{\text{tot}} = 4.1 \qquad f_B^{\text{tot}} = 3.2 \qquad f_{PD}^V = 0.30 \qquad f_{BD}^V = 2.8$$

The lysis percentages reported in the main text correspond to

$$\mathcal{L}^{\text{flux}} = \frac{F_{PD}^V + F_{BD}^V}{F_P^{\text{tot}}} = f_{PD}^V + f_{BD}^V$$

for the cases of weak, strong and very strong recycling. Note that a FN model does not provide information on the carbon stocks $P_c$ and $B_c$, and thus we cannot evaluate $\mathcal{L}_P^{\text{proc}}$ and $\mathcal{L}_B^{\text{proc}}$.

### S5 Mapping models from literature to reference flux network

In this the appendix we describe how the models listed in Table 3 are modified and transformed into a unified model structure. This common structure forms the basis of the PCA analysis presented in Fig. 3.

We build on the minimal microbial ecosystem introduced in Sect. S1. As in Sect. S4, we add a (virtual) compartment $D^V$ through which the lysis flux passes before being transferred to the DOC compartment. This allows us to track the virus-mediated fluxes directly.

For the DS models, we compute the stationary fluxes and normalise them by primary production,

$$f_i^{\text{tot}} = \frac{F_i^{\text{tot}}}{F_P^{\text{tot}}} \qquad \text{and} \qquad f_{ij} = \frac{F_{ij}}{F_P^{\text{tot}}}.$$

These normalised fluxes are then assembled into the flux matrix $F$,

$$F = \begin{pmatrix} 1 & 0 & -f_{PG} & -f_{PD}^0 & -f_{PD}^V \\ 0 & f_B^{\text{tot}} & -f_{BG} & -f_{BD}^0 & -f_{BD}^V \\ 0 & 0 & f_G^{\text{tot}} & -f_{GD} & 0 \\ 0 & -f_{DB} & 0 & f_D^{\text{tot}} & 0 \\ 0 & 0 & 0 & -f_V^{\text{tot}} & f_V^{\text{tot}} \end{pmatrix} \tag{S8}$$

Here we introduced the through-flow flux $f_V^{\text{tot}}$ for the compartment $D^V$, given by

$$f_V^{\text{tot}} = f_{PD}^V + f_{BD}^V,$$

which is entirely transferred to the DOC compartment. Note that the row sums of the flux matrix $F$ give the carbon fluxes lost from the system (through respiration or export) relative to primary production.

For the FN models, we construct the partitioning ratio matrix $A$, solve for the through-flow fluxes using eq. (S6), and normalise them by primary production. From these normalised through-flow and inter-compartment fluxes, we then assemble the corresponding flux matrix $F$ as in eq. (S8).

**Models FUH-B and FUH-PB** are FN models comparable to our minimal microbial ecosystem, except that they include three zooplankton classes (nano-, micro- and nanozooplankton). We merged these into a single grazer compartment $G$ by summing the grazing fluxes entering the three zooplankton compartments and summing the DOC release fluxes leaving them.

Model FUH-B, which includes only viruses infecting heterotrophic bacteria, has the following flux matrix:

$$F_{\text{FUH-B}} = \begin{pmatrix} 1 & 0 & -0.70 & -0.30 & 0 \\ 0 & 0.76 & -0.19 & 0 & -0.19 \\ 0 & 0 & 0.89 & -0.27 & 0 \\ 0 & -0.76 & 0 & 0.76 & 0 \\ 0 & 0 & 0 & -0.19 & 0.19 \end{pmatrix}$$

Model FUH-PB, which includes both viruses infecting phytoplankton and viruses infecting bacteria, has the following flux matrix:

$$F_{\text{FUH-PB}} = \begin{pmatrix} 1 & 0 & -0.63 & -0.34 & -0.03 \\ 0 & 0.80 & -0.20 & 0 & -0.20 \\ 0 & 0 & 0.83 & -0.26 & 0 \\ 0 & -0.80 & 0 & 0.83 & 0 \\ 0 & 0 & 0 & -0.23 & 0.23 \end{pmatrix}$$

**Model WIL** is an FN model similar to FUH-B and FUH-PB. To account for uncertainty, fluxes are given as ranges rather than single values. We sampled values uniformly within these ranges, obtaining 1000 realisations of the model. The average flux matrix across these realisations is given by

$$\overline{F}_{\text{WIL}} = \begin{pmatrix} 1 & 0 & -0.84 & -0.10 & -0.06 \\ 0 & 0.30 & -0.05 & 0 & -0.07 \\ 0 & 0 & 0.89 & -0.30 & -0.01 \\ 0 & -0.30 & 0 & 0.54 & 0 \\ 0 & 0 & 0 & -0.14 & 0.14 \end{pmatrix}$$

Note that this model also includes a (small) viral lysis flux leaving the grazer compartment.

**Model WEI** is the DS model on which the minimal microbial ecosystem model of Sect. S1 is based. Two phytoplankton compartments (cyanobacteria and eukaryotic phytoplankton) were merged and all fluxes were expressed in carbon units rather than nitrogen units, as in the original study. We generated 10000 parameter sets and calculated their corresponding steady states. Using the target compartment densities reported in the original study, we selected the 1000 realisations that were closest to these targets for further analysis.

The average flux matrix across these realisations is given by

$$\overline{F}_{\text{WEI}} = \begin{pmatrix} 1 & 0 & -0.01 & -0.08 & -0.85 \\ 0 & 0.11 & 0 & -0.02 & -0.08 \\ 0 & 0 & 0.02 & 0 & 0 \\ 0 & -0.11 & 0 & 1.03 & 0 \\ 0 & 0 & 0 & -0.93 & 0.93 \end{pmatrix}$$

In this model carbon is strongly recycled within the ecosystem, as indicated by the through-flow of the DOC compartment exceeding primary production, $F_D^{\text{tot}} = 1.03\ F_P^{\text{tot}}$ (averaged across model realisations).

**Models MOJ-S and MOJ-N** are FN models similar to FUH-B and FUH-PB. They were calibrated using data from an oceanographic campaign in the Northeast Atlantic Ocean, corresponding to the southern (MOJ-S) and northern (MOJ-N) regions.

Model MOJ-S has the following flux matrix:

$$F_{\text{MOJ-S}} = \begin{pmatrix} 1 & 0 & -0.25 & -0.20 & -0.35 \\ 0 & 0.85 & -0.36 & -0.04 & -0.45 \\ 0 & 0 & 0.61 & -0.13 & 0 \\ 0 & -0.85 & 0 & 1.17 & 0 \\ 0 & 0 & 0 & -0.80 & 0.80 \end{pmatrix}$$

Model MOJ-N has the following flux matrix:

$$F_{\text{MOJ-N}} = \begin{pmatrix} 1 & 0 & -0.25 & -0.35 & -0.21 \\ 0 & 0.22 & -0.06 & -0.04 & -0.12 \\ 0 & 0 & 0.31 & -0.07 & 0 \\ 0 & -0.22 & 0 & 0.78 & 0 \\ 0 & 0 & 0 & -0.33 & 0.33 \end{pmatrix}$$

Note the strong recycling, quantified by the DOC through-flow relative to primary production, especially for model MOJ-S.

**Models XIE-H and XIE-A** are DS models with detailed spatio-temporal resolution of two contrasted oceanic regions. The XIE-H model is based on the HOT (Hawaii Ocean Time-series) site, an oligotrophic location in the North Pacific Subtropical Gyre with low nutrient concentrations and weak seasonal variability. The XIE-A model corresponds to an Arabian Sea site strongly influenced by seasonal monsoons, resulting in large environmental variability.

We use the carbon fluxes averaged over the year and over the water column, as reported in the Supporting Information of the original study. Model XIE-H has the following flux matrix:

$$F_{\text{XIE-H}} = \begin{pmatrix} 1 & 0 & -0.80 & -0.18 & 0 \\ 0 & 0.52 & -0.12 & -0.11 & -0.01 \\ 0 & 0 & 0.92 & -0.22 & 0 \\ 0 & -0.52 & 0 & 0.52 & 0 \\ 0 & 0 & 0 & -0.01 & 0.01 \end{pmatrix}$$

Model XIE-A has the following flux matrix:

$$F_{\text{XIE-A}} = \begin{pmatrix} 1 & 0 & -0.55 & -0.30 & 0 \\ 0 & 0.70 & -0.17 & -0.17 & -0.01 \\ 0 & 0 & 0.72 & -0.22 & 0 \\ 0 & -0.70 & 0 & 0.70 & 0 \\ 0 & 0 & 0 & -0.01 & 0.01 \end{pmatrix}$$

We performed the PCA using the flux matrices of the different models. A caveat arises, however, because the models differ in how carbon losses associated with bacterial utilisation of DOC are represented. In some models (e.g., WEI and MOJ-N), this loss is placed upstream of the bacterial compartment: a fraction of the DOC through-flow is lost directly from the DOC pool before transfer to bacteria. In other models (e.g., FUH-B, FUH-BP, XIE-H and XIE-A), the loss is represented downstream of the bacterial compartment: DOC uptake increases the bacterial carbon stock, followed by a loss flux from the bacterial compartment itself.

Importantly, these alternative representations of bacterial growth efficiency lead to identical carbon fluxes at the ecosystem level. To enable a robust comparison across models, we therefore introduce aggregate fluxes that combine DOC uptake and subsequent bacterial losses:

$$\begin{aligned} a_{DBG} &= a_{DB}\, a_{BG} = \frac{f_{BG}}{f_D^{\text{tot}}} \\ a^0_{DBD} &= a_{DB}\, a^0_{BD} = \frac{f^0_{BD}}{f_D^{\text{tot}}} \\ a^V_{DBD} &= a_{DB}\, a^V_{BD} = \frac{f^V_{BD}}{f_D^{\text{tot}}} \end{aligned}$$

The PCA analysis was performed using the following ten variables:

| | |
|---|---|
| $f_G^{\text{tot}}$ | through-flow of $G$ compartment relative to primary production |
| $f_D^{\text{tot}}$ | through-flow of $D$ compartment relative to primary production |
| $f_V^{\text{tot}}$ | total viral lysis flux relative to primary production |
| $a_{PG} = \dfrac{f^V_{PG}}{f_P^{\text{tot}}}$ | grazing partitioning ratio of phytoplankton |
| $a^0_{PD} = \dfrac{f^0_{PD}}{f_P^{\text{tot}}}$ | DOC partitioning ratio of phytoplankton (excluding viral lysis) |

$$a^V_{PD} = \frac{f^V_{PD}}{f^{\text{tot}}_P} \quad \text{viral lysis partitioning ratio of phytoplankton}$$

$$a_{DBG} = \frac{f_{BG}}{f^{\text{tot}}_D} \quad \text{grazing partitioning ratio of bacteria}$$

$$a^0_{DBD} = \frac{f^0_{BD}}{f^{\text{tot}}_D} \quad \text{DOC partitioning ratio of bacteria (excluding viral lysis)}$$

$$a^V_{DBD} = \frac{f^V_{BD}}{f^{\text{tot}}_D} \quad \text{viral lysis partitioning ratio of bacteria}$$

$$a_{GD} = \frac{f_{GD}}{f^{\text{tot}}_G} \quad \text{DOC partitioning ratio of grazer compartment}$$

The histograms of these variables for the different models are shown in Fig. S1.

To ensure equal weighting of all models in the PCA, a balanced sampling strategy was applied, such that each model contributed the same total number of realisations. Models WIL and WEI, which incorporate parameter uncertainty, were represented by 1000 distinct model realisations each. For the FUH, MOJ and XIE models, which each comprise two realisations, 500 identical copies of each realisation were included. As a result, each model contributed a total of 1000 realisations to the PCA, ensuring equal overall influence on the analysis.

### S6 Studying virus removal: technical details

To study the effects of virus removal, we use the minimal microbial ecosystem model introduced in Sect. S1, rather than the more elaborate model of Weitz *et al.* (2015). In the latter, cyanobacteria $C$ and eukaryotic phytoplankton $E$ are dealt with separately, each controlled by its own virus population, $V_C$ and $V_E$. In such a setting, removing viruses inevitably leads to the extinction of one phytoplankton group. Because here we focus on the virus impact on the carbon flux network, rather than on virus-mediated coexistence, we prefer to work with a model with a single phytoplankton compartment, which persistence does not depend on viruses.

For the first analysis (Fig. 4), we sampled parameter values from Table S2, and selected those for which the equilibrium state lay within the ranges specified in Table S1. We then removed viruses from the model and simulated the new stationary state in their absence. Out of the 10000 parameter sets with viruses, 8651 reached equilibrium after virus removal. We constructed the flux matrices for the system with and without viruses. From these matrices, we computed the absolute and relative viral lysis fluxes ($F^{\text{tot}}_V$, $f^{\text{tot}}_V = F^{\text{tot}}_V / F^{\text{tot}}_P$ and $F^{\text{tot}}_V / F^{\text{tot}}_D$), as well as the absolute and relative changes induced by virus removal. We quantified these changes for primary production ($\Delta F^{\text{tot}}_P$ and $\delta F^{\text{tot}}_P$) and for DOC though-flow ($\Delta F^{\text{tot}}_D$ and $\delta F^{\text{tot}}_D$). Most of these variables have strongly skewed distribution (Fig. S2), which could bias correlation plots. We therefore applied a rank-based normal transformation (sometimes called gaussianisation), which maps values onto a Gaussian distribution while preserving their relative ordering. Fig. S3 shows the correlation plot for all variable pairs, while Fig. 4 in the main text shows a subset of these pairs.

For the second analysis (Fig. 5), in which we compare the effects of virus removal between the DS and FN approaches, we proceeded initially as in the first analysis. In addition to removing viruses in the DS models, we also removed viruses using the FN approach. To do this, we set the viral fluxes

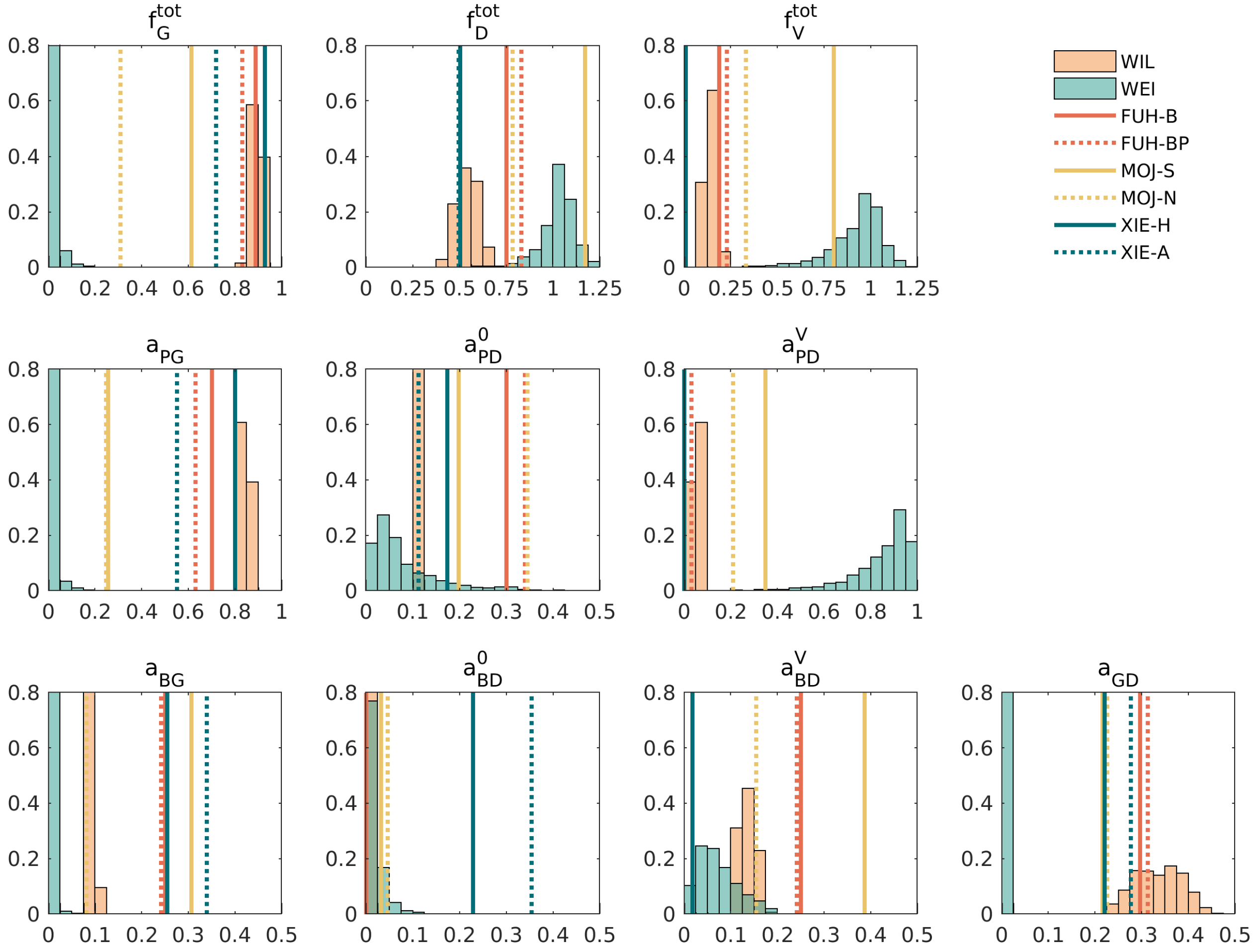


Figure S1: Histogram of the ecosystem variables used in the PCA analysis. The distribution of the 1000 realisations of WIL and WEI models is represented as histograms. For models without parameter variability, results are displayed as vertical lines. Solid and dashed lines correspond to different parameterisations of the same model.

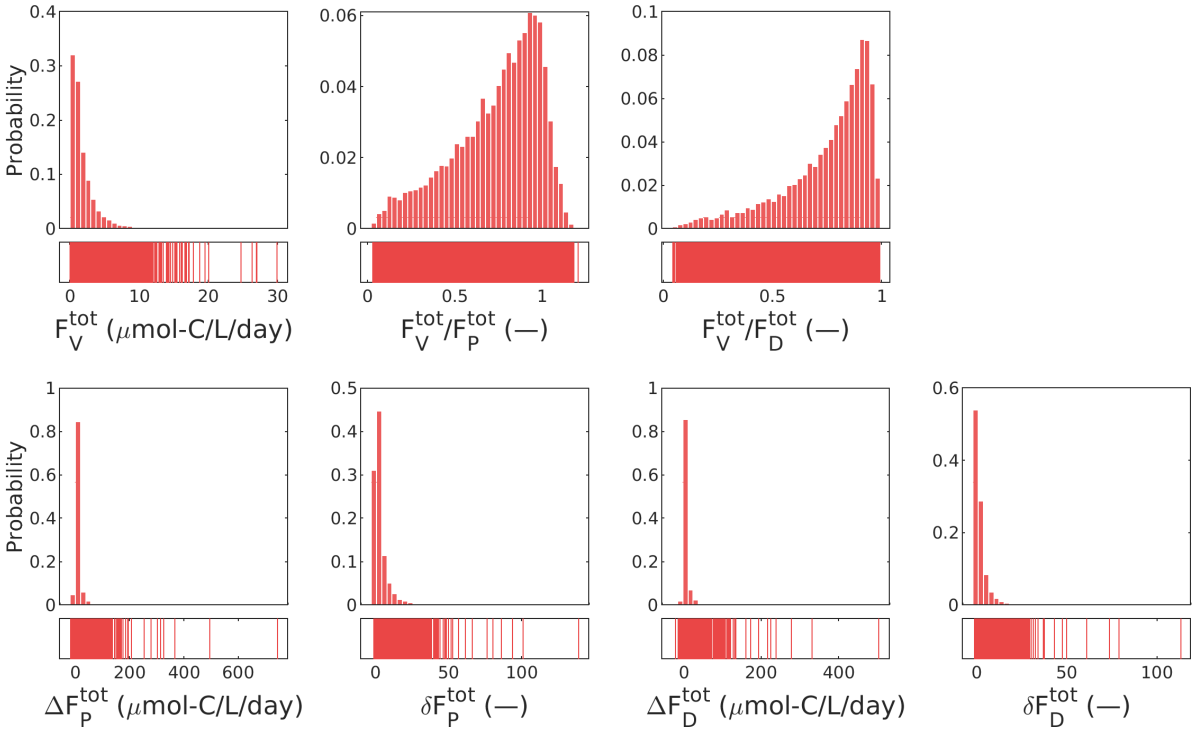


Figure S2: Distributions of variables from the study of virus removal impacts. Top row: viral lysis fluxes. Bottom row: changes induced by virus removal. Each variable is represented by two panels: the top panel shows a histogram and the bottom panel shows individual values as vertical lines to highlight outliers that may not be visible in the histogram. Most variables have a strongly skewed distribution.

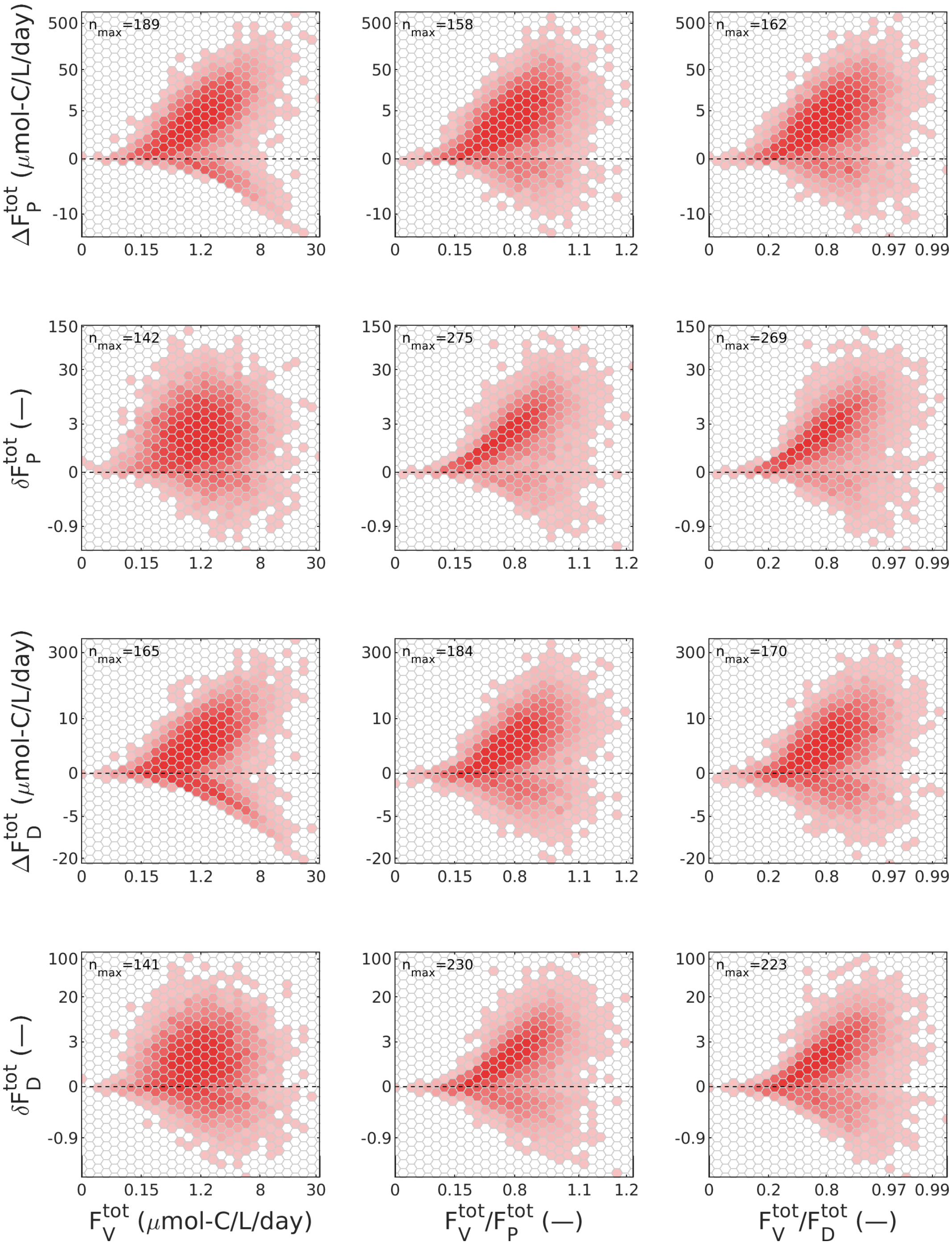


Figure S3: Same as Fig. 4 but for an extended set of variables. On the $x$-axes: absolute viral lysis flux $F_V^{\text{tot}}$; viral lysis flux relative to primary production $F_V^{\text{tot}}/F_P^{\text{tot}}$ and relative to the DOC through-flow $F_V^{\text{tot}}/F_D^{\text{tot}}$. On the $y$-axes: absolute and relative change in primary production, $\Delta F_P^{\text{tot}}$ and $\delta F_P^{\text{tot}}$; absolute and relative change in DOC through-flow, $\Delta F_D^{\text{tot}}$ and $\delta F_D^{\text{tot}}$.

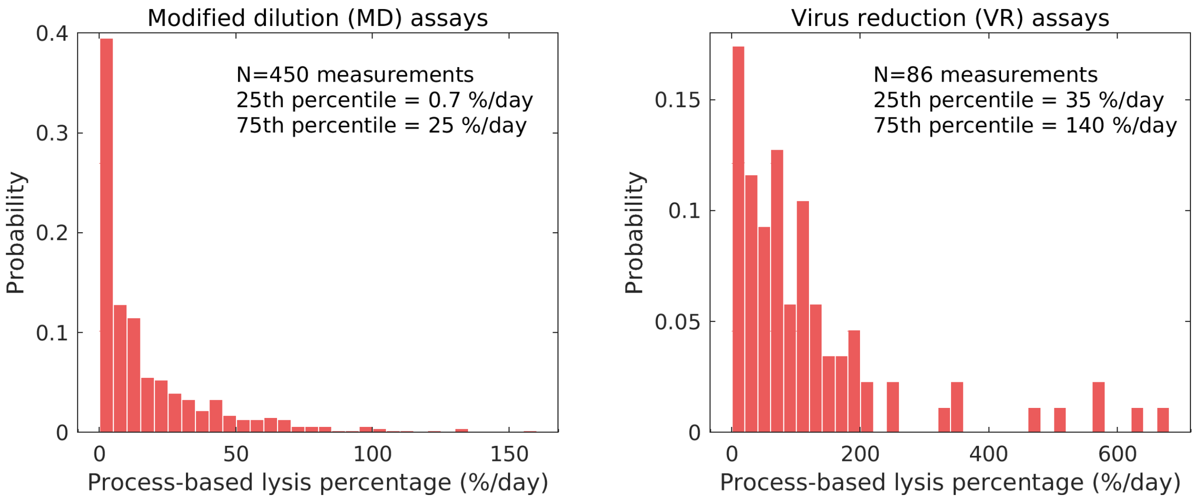


Figure S4: Distributions of experimentally estimated viral lysis rates compiled from Mojica & Brussaard (2026). Left panel: lysis rates of phytoplankton from MD assays. Right panel: lysis rates of heterotrophic bacteria from VR assays.

to zero and rescaled the remaining flux partitioning ratios. For example, for the phytoplankton compartment, if the partitioning ratios before virus removal were

$$a_{PG},\ a^0_{PD},\ a^V_{PD},\quad \text{with a loss fraction } a_{P0} = 1 - a_{PG} - a^0_{PD} - a^V_{PD}$$

then after virus removal the rescaled ratios were

$$\frac{a_{PG}}{a_{PG} + a^0_{PD} + a_{P0}},\ \frac{a^0_{PD}}{a_{PG} + a^0_{PD} + a_{P0}},\ 0,\quad \text{and a loss fraction } \frac{a_{P0}}{a_{PG} + a^0_{PD} + a_{P0}}$$

which can also be written as

$$\frac{a_{PG}}{1 - a^V_{PD}},\ \frac{a^0_{PD}}{1 - a^V_{PD}},\ 0,\quad \text{and a loss fraction } \frac{a_{P0}}{1 - a^V_{PD}}.$$

We then computed the stationary fluxes for these modified partitioning ratios using eq. (S6). Selected through-flow fluxes and partitioning ratios after virus removal were compared between the DS and FN approaches.

### S7 Ranges of experimental viral lysis rates

We extracted the experimental data from the supplementary files accompanying Mojica & Brussaard (2026). For the modified dilution (MD) assays, we used the column "Viral lysis rate ($d^{-1}$)" from the file `MDA_data.xlsx`. For the virus reduction (VR) assays, we used the columns "VP ($ml^{-1}$ $d^{-1}$)", "Bacteria ($ml^{-1}$)", "BS (viruses $cell^{-1}$)" from the file `VP_data.xlsx`, and computed lysis rates by dividing viral production by bacterial abundance and burst size. The resulting distributions are shown in Fig. S4.
